# Bioorthogonal labeling of transmembrane proteins with non-canonical amino acids allows access to masked epitopes in live neurons

**DOI:** 10.1101/2021.02.27.433189

**Authors:** Diogo Bessa-Neto, Alexander Kuhlemann, Gerti Beliu, Valeria Pecoraro, Sören Doose, Natacha Retailleau, Nicolas Chevrier, David Perrais, Markus Sauer, Daniel Choquet

## Abstract

Progress in biological imaging is intrinsically linked to advances in labeling methods. The explosion in the development of high-resolution and super-resolution imaging calls for new approaches to label targets with small probes. These should allow to faithfully report the localization of the target within the imaging resolution – typically nowadays a few nanometers - and allow access to any epitope of the target, in the native cellular and tissue environment. We report here the development of a complete labeling and imaging pipeline using genetic code expansion and non-canonical amino acids in primary neurons that allows to fluorescently label masked epitopes in target transmembrane proteins in live neurons, both in dissociated culture and organotypic brain slices. This allowed us to image the differential localization of two glutamate receptor auxiliary proteins in complex with their partner with a variety of methods including widefield, confocal, and *d*STORM super-resolution microscopy.

## INTRODUCTION

Over the past 15 years, advances in light-based super-resolution microscopy have revolutionized the way neuroscientists perceive key neuronal processes such as synaptic and axonal nanoscale organization or protein trafficking at the single-molecule level^1–3^. The improvements in the various super-resolution imaging methods, and particularly in single-molecule localization microscopy (SMLM), have made it possible to routinely reach imaging resolutions in the order of ~20 nanometers. However, elucidating target protein organization at virtually molecular resolution requires not only a high localization precision of individual emitters but also a high labeling density and specificity, and a distance between the fluorescent reporter and the target (linkage error) substantially smaller than the desired imaging resolution^4,5^.

In general, two broad categories of labeling methods are used for fluorescence imaging: either labeling the target protein with a ligand-dye complex or genetic fusion with a reporter fluorescent protein. In the first case, the oldest and most widespread approach is immunostaining using a complex of primary and secondary fluorescently labeled antibodies. This results in linkage errors of typically 10-20 nm as whole-IgG antibodies (Abs) are large proteins (10-15 nm)^6^, limiting the resolution achievable even with optimized super-resolution microscopy^7^. Another pitfall of using Abs in live-cell experiments is their multivalence, which may induce protein crosslinking and, therefore, modify target function or create artificial clusters. Over the years, new smaller and monovalent alternatives to label endogenous proteins have emerged such as ligands derived from antibodies (Fab, scFv, VhH), affimers, or aptamers^8–12^. However, the availability of such compounds for many targets is still limited and the linkage error remains at least 4 nm for the smallest ones. In the alternative labeling method, gene fusion technology with fluorescent proteins which are suitable for super-resolution microscopy, has been a reliable tool due to its versatility, stoichiometric labeling of the target, and specificity. However, these protein tags still result in a complex in the order of 5 nm. In addition, while considerably smaller than a whole-IgG Ab, the incorporation of such bulky components to a target protein can impede their native function, accounting for biased interpretations and severely limiting their possible site of insertion.

Click chemistry labeling via genetic code expansion (GCE) offers the possibility for site-specific incorporation of non-canonical amino acids (ncAAs) containing bioorthogonal groups into a target protein^13,14^. By replacing a native codon at a selected position in the target protein with a rare codon, such as the Amber (TAG) stop codon, the modified protein can then be expressed into the desired host cells along with an engineered amino acyltransferase (RS) and tRNA pair orthogonal to the host translational machinery. The engineered RS is modified in a way to only recognize a specific ncAA, which is then attached to a tRNA that matches the rare codon. Among a collection of different possibilities, the trans-cyclooct-2-ene (TCO*)-modified amino acids, as TCO*-L-lysine (TCO*-A), is of interest when it comes to targeting and labeling the desired target proteins in living organisms. TCO* can react with a 1,2,4,5-tetrazine in a catalyst-free, fast, specific, and bioorthogonal strain-promoted inverse electron-demand Diels-Alder cycloaddition reaction (SPIEDAC). Due to the high selectivity and fast kinetics of this click chemistry reaction, a large number of fluorophore-tetrazine conjugates and TCO* functionalized molecules are now commercially available making labeling of mammalian cells and whole organisms with organic dyes accessible for live and fixed samples,^15–17^.

The nanoscale organization of synapses is an ideal model system for the application of innovative imaging and labeling methods because of its exquisite complexity and diversity as well as because of the tight link between synapse dynamic organization and function^2^. Among synaptic proteins, the complex involved in regulation of the function, localization, and trafficking of AMPA receptors (AMPAR) – the glutamate receptors that mediate most excitatory synaptic transmission, has historically raised large interest. Transmembrane AMPAR regulatory protein (TARP) family are four transmembrane proteins characterized by an intracellular amino- and carboxyl-terminal domain, and two extracellular loops (Ex1 and Ex2)^18^ (Supplementary Fig. 1a). TARPs are key modulators of AMPAR-mediated synaptic transmission and plasticity, as they promote surface targeting, AMPAR pharmacology and gating modulation fundamental for proper AMPAR-mediated transmission^19–25^. Among the different members of the TARP family, γ2 (also known as stargazin) is the prototypical AMPAR-auxiliary protein and has been the most widely studied, followed, more recently, by γ8 that is the most abundant TARP in the hippocampus^26^. While sharing a large homology^27^, γ2 and γ8 not only exert a differential AMPAR-modulation^25, 28, 29^ but also display differential plasma membrane distribution, with γ2 suggested to bear an almost exclusive synaptic localization and γ8 a more widespread dendritic distribution as seen in electron microscopy studies^30,31^. This was never, to the best of our knowledge, confirmed in living neurons by optical microscopy due to the lack of adequate tools. Here, we explored the potential of bioorthogonal labeling as a strategy to tag and visualize surface TARPs in living neurons with minimal to non-perturbation of TARPs modulation over AMPAR, opening new doors to the study of hard-to-tag proteins in living neurons.

## Results

### Epitope masking by close interaction of TARPs extracellular loops with AMPAR LBD

Our understanding of TARPs localization and trafficking has been hampered by a lack of suitable labeling and imaging tools. The close association of the extracellular domains of TARPs to the AMPAR ligand-binding domain (LBD)^32–35^ that confer their role in TARP-specific AMPAR modulation^25, 28, 36, 37^ has hindered the development of ligands recognizing the extracellular domains of TARPs as well as genetic fusion tagging^8,38^. Deciphering the respective surface diffusion and synaptic organization properties of γ2 and γ8 is of particular interest given their presumptive key role in AMPAR regulation and modulation as well as the control of synaptic plasticity, but unfortunately remains still widely unknown.

As a first attempt to create specific ligands for γ2 and γ8 that could be used to study their organization and trafficking in live neurons, we generated antibodies against the γ2 Ex2 and γ8 Ex1 extracellular loops (Supplementary Fig. 1b). We first evaluated the Abs specificity by incubating living COS-7 cells expressing either γ2 or γ8 bearing mEos2 as a reporter. As shown in Supplementary Fig. 1c, both Abs are specific towards their respective target protein. We then analyzed if we could use these Abs to label endogenous TARPs in dissociated hippocampal neurons, as both γ2 and γ8 are expressed in the hippocampus^27, 39, 40^. To our surprise, our Abs were unable to recognize endogenous γ2 or γ8 in our primary hippocampal cultures. The presence of γ8 was confirmed by post-fixation immunostaining against the intracellular C-terminal domain of γ8 (Supplementary Fig. 1d), while γ2 immunostaining was inconclusive due to the poor sensitivity of the tested commercial α-γ2 Abs. We know, however, that γ2 is expressed in hippocampal cultures from biochemistry data (data not shown). We then overexpressed γ2 bearing mCherry (γ2::mCherry) or N-terminal eGFP-tagged GluA2 tethered to γ2 (eGFP::GluA2::γ2), where the GluA2 C-terminus is fused to the γ2 N-terminus by in-frame expression^41^. When labeled with the α-γ2 Ex2, neurons overexpressing γ2::mCherry displayed specific Ab labeling that colocalized with the mCherry signal. In contrast, in neurons overexpressing eGFP::GluA2::γ2 and labeled with the α-γ2 Ex2, no Ab labeling was observed (Supplementary Fig. 1e). Structural data of γ2 or γ8 in complex with AMPARs reveal close proximity of both Ex1 and Ex2 to the AMPAR ligand-binding domain (LBD)^33–35^, which likely results in epitope masking. This finding leads to the conclusion that Abs cannot recognize endogenous TARPs in dissociated hippocampal neurons as well as in eGFP::GluA2::γ2-overexpressing neurons because of steric inaccessibility. Altogether, our results indicate that, at endogenous levels at the plasma membrane, TARPs are always associated with AMPAR that mask the extracellular epitopes.

### Genetic code expansion and bioorthogonal labeling of TARPs

Bioorthogonal labeling of proteins by replacing a single natural amino acid with a non-canonical/unnatural amino acid (ncAA) has emerged in the past years as an alternative strategy to target and visualize proteins in living mammalian cells with minimal to no-perturbation^15,16,42^. Because click chemistry labeling of ncAAs with tetrazine-dyes is efficient and sterically minimally demanding, we hypothesized that it might be the method of choice to label masked epitopes on TARPs. We first designed Amber mutants (herein termed ncAA-tagged) of γ2 and γ8 by site-direct mutagenesis (Fig. 1a), with respect to previously conducted work on γ2 with ncAAs^32^. Additionally, we replaced the endogenous Amber termination codon of γ8 with the Ochre codon (TAA) to prevent erroneous ncAA incorporation at the C-terminus (see Methods section). To identify the best position for the Amber mutation, three mutants of each, γ2 (S44*, S51* and S61*) and γ8 (S72*, S84* and K102*) were tested for ncAA incorporation by expression level and labeling efficiency in HEK293T cells (Fig. 1b-d and Supplementary Fig. 1f,g). To check the efficiency of ncAA incorporation in the different mutants, eGFP was fused within the C-tail of γ2 and γ8, downstream the Amber mutation. Hence, inefficient incorporation or the absence of ncAAs would result in premature translation termination and loss of eGFP signal (Fig. 1c,d). HEK293T cells were co-transfected with the different ncAA-tagged TARPs and an engineered pyrrolysine aminoacyl tRNA synthetase and its cognate tRNA, respectively, PylRS, and single-copy tRNA^Pyl^ (tRNA^Pyl^)^43^ or four copies (4xtRNA^Pyl^)^44^. The clickable trans-cyclooctene derivatized lysine (TCO*A) was added to the cell media at the time of transfection for a period of approximately 24 h. Surface expression of the ncAA-tagged TARPs was accessed by bioorthogonal labeling via SPIEDACusing cell-impermeable tetrazine-dyes (H-Tet-Cy3, H-Tet-Cy5 and Pyr-Tet-ATTO643), which react with TCO* in a catalysis-free ‘click-reaction’^15,16,45^. Afterward, excess of tetrazine-dyes was removed by subsequent washes and cells were live-imaged using confocal microscopy. As indicated by the eGFP signal, all ncAA-tagged TARPs showed comparable expression levels as compared to γ2::eGFP, revealing efficient incorporation of TCO*A. However, the mutant γ8 K102*::eGFP displayed decreased Pyr-Tet-ATTO643 labeling as compared to mutants γ8 S72*::eGFP and γ8 S84*::eGFP. No noticeable difference in tetrazine-dye labeling efficiency was observed within the different γ2 mutants (Supplementary Fig. 1f,g). Additionally, tetrazine-dye labeling was entirely due to the incorporation of TCO*A as no labeling was detected in cells transfected with γ2::eGFP in the presence of TCO*A nor in cells transfected with γ2 S44*::eGFP or γ8 S72*::eGFP in the absence of TCO*A (Fig. 1c,d and Supplementary Fig. 1f).

**Figure 1:**
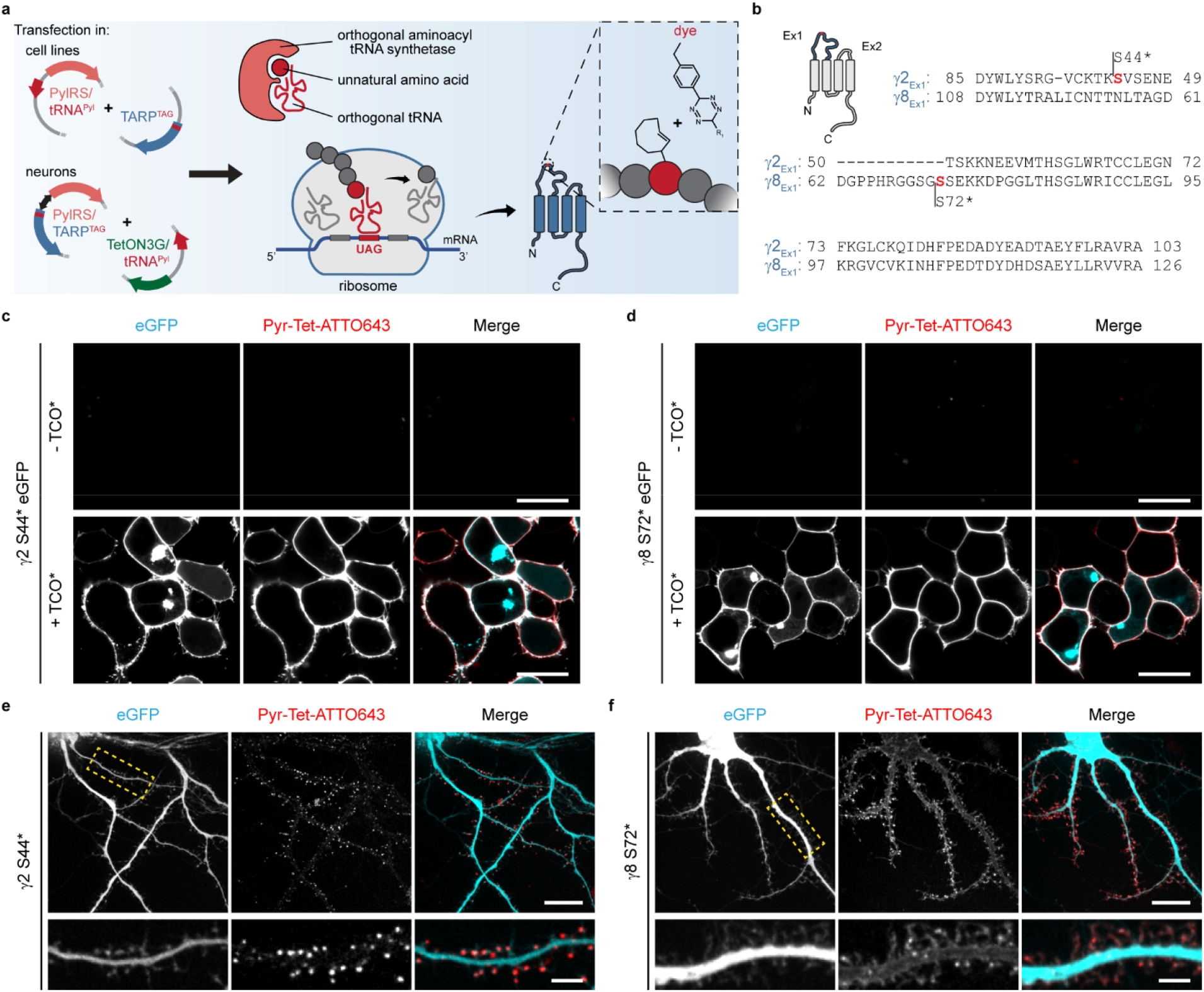
Bioorthogonal labeling of TARPs. (**a**) Schematic overview of click chemistry labeling via genetic code expansion. Amber mutants of ncAA-tagged γ2 and γ8 were designed by standard site-direct mutagenesis to incorporate the ncAAs carrying a TCO*A for click labeling. Protein expression occurs through endogenous and orthogonal tRNA-synthetases in the presence of appropriate tRNAs. Labeling of ncAA-tagged proteins occurs through a catalysis-free reaction between TCO*A and tetrazine-functionalized dyes via the strain-promoted inverse electrondemanding Diels-Alder cycloaddition (SPIEDAC) reaction. (**b**) Sequence alignment of the first extracellular loops of γ2 and γ8 from *rattus norvegicus.* Amber substitution mutations are represented in red. (**c-d**) Representative confocal images of living HEK293T cells co-expressing PylRS/4xtRNA^Pyl^ and (**c**) γ2 S44*::eGFP or (**d**) γ8 S72*::eGFP in the absence (upper) or presence of 250 μM TCO*A (lower) stained with 1.5 μM Pyr-Tet-ATTO643. Scale bar: 20 μm. (**e-f**) Representative spinning disk confocal images of living dissociated hippocampal neurons coexpressing eGFP, Tet3G/tRNA^Pyl^ and (**e**) pTRE3G-BI PylRS/γ2 S44* or (**f**) pTRE3G-BI PylRS/γ8 S72* in the presence of 250 μM TCO*A and 100 ng.mL^-1^ doxycycline labeled with 0.5 μM Pyr-Tet-ATTO643. Lower panels, magnified views of segments of dendrites highlighted in the eGFP channels (dashed yellow boxes) of the overview images showing the distribution of γ2 S44* and γ8 S72*. Scale bar: 20 μm (overview images) and 5 μm (magnified images).

### ncAA-tagged TARPs physically and functionally interact with AMPAR-subunit GluA1 as seen by FRET and electrophysiology

TARPs are auxiliary subunits to AMPARs that bind and interact closely with GluA subunits, as demonstrated by biochemical^23,38^, functional^23^ and structural^33–35^ data. We thus aimed to study whether ncAA-tagged TARPs could still physically and functionally interact with GluA subunits using both Förster Resonance Energy Transfer (FRET) measurements and electrophysiology. As we were able to insert ncAAs into the Ex1 loops of γ2 and γ8 that are in close apposition to the extracellular domain of GluA subunits^33–35^, we first sought to use FRET to measure AMPAR-TARP interactions. We designed a set of possible FRET pairs between the ncAA-tagged γ2/γ8 and the AMPAR-subunit GluA1. To label surface GluA1, a SNAP-tag was either inserted at the N-terminus (SNAP::GluA1; no FRET expected) or within the ATD-LBD linker of GluA1 at position 396 aa (GluA1::SNAP396; potential FRET pair) (Fig. 2a). HEK293T cells were co-transfected with PylRS/tRNA^Pyl^, SNAP-tagged GluA1 and ncAA-tagged TARPs in the presence of TCO*A. Cells were stained with 5 μM BG-AF488 (donor) and 1.5 μM H-Tet-Cy3 (acceptor) at 37 °C for 30 minutes. Excess dye was subsequently removed by washing steps with HBSS. Fluorescence lifetime imaging microscopy (FLIM) was used to estimate the degree of FRET-based changes of the donor AF488 fluorescence lifetime^46^. As expected, when co-expressed with the SNAP::GluA1-AF488, neither γ2 S44*-Cy3 nor γ8 S72*-Cy3 were able to quench the donor, therefore, no decrease in fluorescence lifetime was observed as compared to the donor alone (no H-Tet-Cy3), GluA1::SNAP396-AF488 + γ2 S44* (data not shown) (SNAP::GluA1-AF488 + γ2 S44*-Cy3: τ_AF488_ = 3.07 ± 0.03 ns; SNAP::GluA1-AF488 + γ8 S72*-Cy3: τ_AF488_ = 3.04 ± 0.01 ns; GluA1::SNAP396-AF488 + γ2 S44*: τ_AF488_ = 3.01 ± 0.01 ns, respectively). In contrast, we observed a robust decrease in GluA1::SNAP396-AF488 fluorescence lifetime when co-expressed with γ2 S44*-Cy3 or γ2 S61*-Cy3 as compared to the SNAP::GluA1-AF488 + γ2 S44*-Cy3, with the FRET pair GluA1::SNAP396-AF488 + γ2 S44*-Cy3 showing a stronger reduction in AF488 lifetime (Fig. 2b,c). Moreover, when we forced a one to one interaction between GluA1 and γ2 using a tethered GluA1 SNAP396 to γ2 S61* (GluA1::SNAP396::γ2 S61*-AF488/Cy3), we did not observe a significant difference compared to the condition in which we expressed the two proteins separately (GluA1::SNAP396-AF488 + γ2 S61*-Cy3). This suggests a full occupancy of the AMPAR subunits with four TARPs under our experimental conditions. Similar to ncAA-tagged γ2-Cy3, the presence of ncAA-tagged γ8-Cy3 led to a robust decrease in GluA1::SNAP396-AF488 lifetime, with the GluA1::SNAP396-AF488 + γ8 S84*-Cy3 pair outperforming the pairs GluA1::SNAP396-AF488 + γ8 S72*- and γ8 K102*-Cy3 (Fig. 2d). As for γ2, we did not observe a difference between the GluA1::SNAP396-AF488 + γ8 K102*-Cy3 and the tethered GluA1 SNAP396 to γ8 K102* (GluA1::SNAP396::γ8 K102*-AF488/Cy3). Altogether, our FRET experiments indicate that ncAA-tagged TARPs physically interact with AMPAR and provide thus a new tool to study the regulation of AMPAR-TARP interactions.

**Figure 2:**
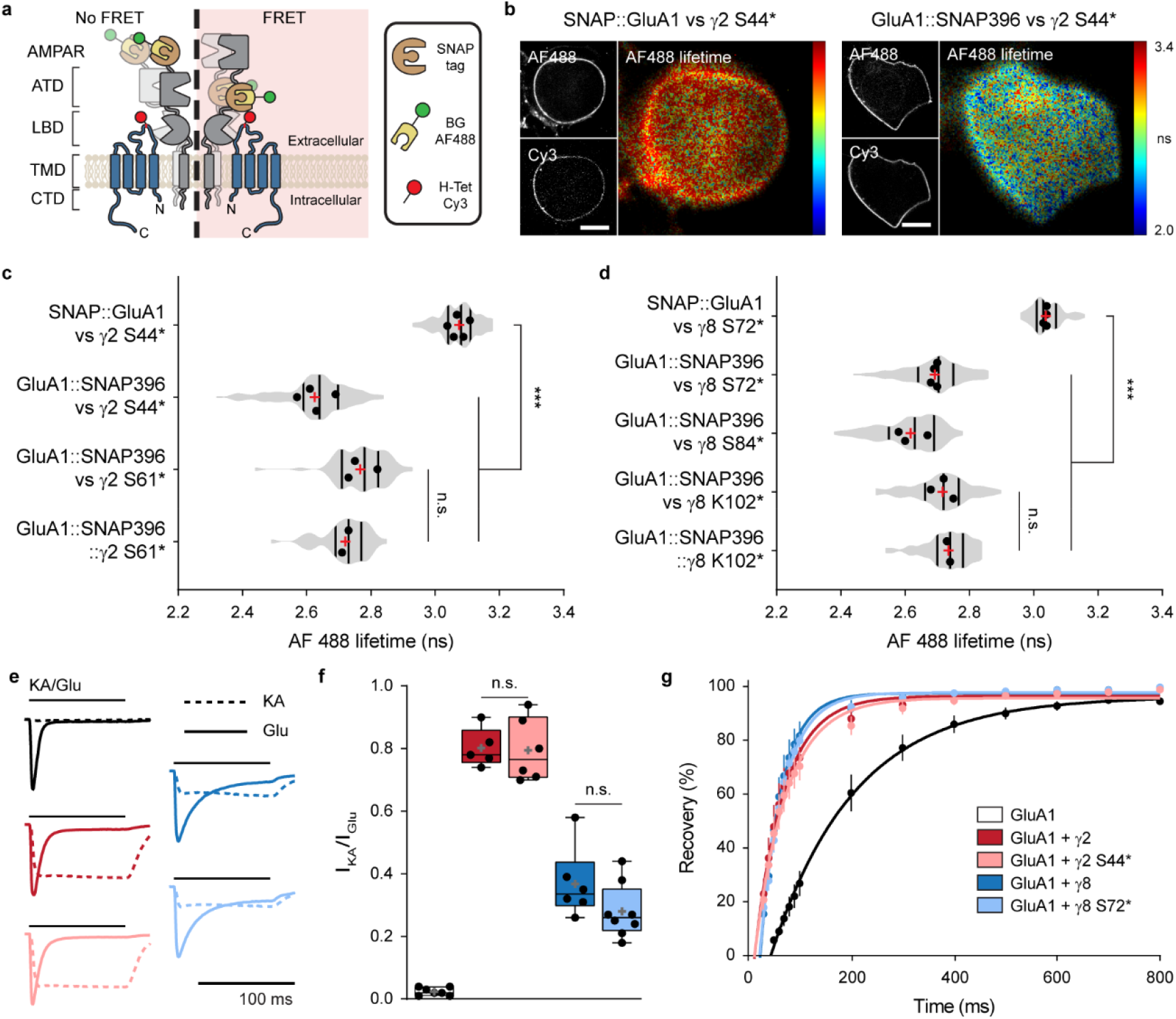
ncAA incorporation in the EX1 loop of TARPs does not impair physical or functional interaction with AMPAR-subunit GluA1. (**a**) Depiction of the strategy used to label AMPAR-subunit GluA1 and TARPs for Förster resonance energy transfer (FRET) microscopy application. As a control, SNAP-tag was inserted at the N-terminus of GluA1, and as a possible FRET pair, SNAP-tag was inserted between the ATD-LBD linker of GluA1, in close proximity to the EX1 loop of TARPs, which was tagged by ncAA incorporation and labeled with tetrazine Cy3 (acceptor). SNAP-tag was labeled with BG-AF488 (donor). (**b**) Representative spinning disk confocal (AF488 and Cy3) and widefield illumination fluorescence lifetime imaging microscopy (FLIM; AF488 lifetime) images of living HEK293T cells co-expressing PylRS/tRNA^Pyl^, γ2 S44*, and SNAP GluA1 (left) or GluA1 SNAP396 (right) stained with 5 μM SNAP-AF488 and 1.5 μM H-Tet-Cy3. Scale bar: 10 μm. (**c**) Average AF488 lifetime measured by FLIM in HEK293T cells coexpressing PylRS/tRNA^P^y^l^, SNAP::GluA1-AF488 + γ2 S44*-Cy3 (τAF488 = 3.07 ± 0.03 ns; n = 117), GluA1::SNAP396-AF488 + γ2 S44*-Cy3 (τAF488 = 2.63 ± 0.05 ns; n = 108), GluA1::SNAP396-AF488 + γ2 S61*-Cy3 (τ_AF488_ = 2.77 ± 0.05 ns; n = 66), or tethered GluA1::SNAP396::γ2 S61*-AF488/Cy3 (τ_AF488_ = 2.72 ± 0.01 ns; n = 51). (**d**) Average AF488 lifetime measured by FLIM in HEK293T cells co-expressing PylRS/tRNA^Pyl^, SNAP::GluA1-AF488 + γ8 S72*-Cy3 (τ_AF488_ = 3.04 ± 0.01 ns; n = 87), GluA1::SNAP396-AF488 + γ8 S72*-Cy3 (τ_AF488_ = 2.69 ± 0.01 ns; n = 104), GluA1::SNAP396-AF488 + γ8 S84*-Cy3 (τ_AF488_ = 2.62 ± 0.04 ns; n = 111), GluA1::SNAP396-AF488 + γ8 K102*-Cy3 (2.72 ± 0.04 ns; n = 84), or tethered GluA1::SNAP396::γ2 K102*-AF488/Cy3 (τ_AF488_ = 2.74 ± 0.01 ns; n = 53). (**e-f**) Representative whole-cell currents (**e**) and (**f**) ratios of kainate (KA)- to glutamate (Glu)-evoked currents in response to 0.1 mM kainate (KA; dashed) or 10 mM glutamate (Glu; line) in GluA1- and PylRS/tRNA^Pyl^-positive HEK293T cells coexpressing eGFP (control; 0.02 ± 0.01; n= 7; black), γ2::eGFP (0.80 ± 0.06; n = 5; red), γ2 S44*::eGFP (0.80 ± 0.10; n = 6; light red), γ8::eGFP (0.37 ± 0.11; n = 6; blue), or γ8 S72*::eGFP (0.28 ± 0.09; n = 8; light blue). (**g**) Recovery of desensitization to two pulses of 100 ms Glu applied at different intervals to whole-cell patches from HEK293T cells co-expressing PylRS/tRNA^Pyl^, GluA1 and, eGFP (τ_rec_ = 162.6 ms; n = 6; black), γ2::eGFP (τ_rec_ = 57.1 ms; n = 5; blue), γ2 S44*::eGFP (τ_rec_ = 62.3 ms; n = 6; light blue), γ8::eGFP (τ_rec_ = 40.9 ms; n = 6; red), and γ8 S72*::eGFP (τrec = 45.8 ms; n = 8; light red). All data represent mean ± standard deviation.

TARPs subtype I, including γ2 and γ8, modulate AMPAR gating in a TARP subtype-specific manner^25,28,29,36,47^. To determine if the incorporation of TCO*A within the Ex1 loop compromises TARP function, notably interaction with and modulation of AMPARs, we performed whole-cell patch-clamp recordings in HEK293T cells co-expressing GluA1 (we used the flip isoform) alone (eGFP, control) or in the presence of WT or ncAA-tagged γ2/γ8, bearing eGFP as a reporter. When compared to GluA1 alone and GluA1 plus respective WT TARP, the incorporation of TCO*A into the EX1 loop did not impair the ability of TARPs to increase the efficacy of the partial agonist kainate (KA) over GluA1 (ratio peak current amplitude KA/Glu mean ± SD: GluA1 = 0.02 ± 0.01; GluA1 + γ2 = 0.80 ± 0.06; GluA1 + γ2 S44* = 0.80 ± 0.10; GluA1 + γ8 = 0.37 ± 0.11; GluA1 + γ8 S72* = 0.28 ± 0.09) (Fig. 2e,f). Additionally, we did not observe perturbations on TARPs ability to decrease receptor desensitization (τ_des_ in ms: GluA1 = 4.34 ± 0.59; GluA1 + γ2 = 8.44 ± 1.50; GluA1 + γ2 S44* = 9.02 ± 2.14; GluA1 + γ8 = 17.41 ± 3.72; GluA1 + γ8 S72* = 15.43 ± 2.82) (Supplementary Fig. 2a) or increase receptor recovery from desensitization (τ_rec_, in ms: GluA1 = 162.6; GluA1 + γ2 = 57.1; GluA1 + γ2 S44* = 62.3; GluA1 + γ8 = 40.9; GluA1 + γ8 S72* = 45.8) (Fig. 2g). Furthermore, no difference was observed between WT TARP and respective ncAA-tagged TARP ability to reduce the inwardly rectified currents observed in GluA1 in the absence of TARPs (Supplementary Fig. 2b). Altogether, the patch-clamp experiments indicate that ncAA-tagged TARPs retain a normal functional ability to modulate AMPAR gating.

### Distinct surface distributions of ncAA-tagged γ2 and γ8 in hippocampal neurons

The occurrence of the Amber codon in mammalian cells is rare, (~0.5 ‰), and represents 20-23% of all stop codons. It is however important to keep in mind that GCE might induce toxicity due to tRNA modifications of endogenous proteins containing Amber codon terminations. A good practice is to restrict the concentration of PylRS to prevent suppression of naturally occurring Amber codon terminations, as in the CACNG8 gene, encoding the protein γ8. In addition, in our hands, long-term overexpression of TARPs tends to induce toxicity in neurons. Hence, we decided to overexpress the PylRS and ncAA-tagged TARPs under a bidirectional doxycycline-inducible promoter, pTRE3G-BI, i.e. pTRE3G-BI PylRS/γ2 S44* and pTRE3G-BI PylRS/γ8 S72*. To further decrease the complexity of our tool, we combined in a single vector the tRNA^Pyl^ and the Tet-On 3G transactivator (herein termed Tet3G/tRNA^Pyl^), which binds to and activates expression from TRE3G promoters in the presence of doxycycline (see Methods section). Dissociated hippocampal neurons at days *in vitro* (DIV) 3-4 were transfected with the necessary machinery for the expression of γ2 S44* or γ8 S72* together with eGFP. At DIV 16-17, 100 ng mL^-1^ doxycycline and 250 μM TCO*A were added to the cell media for approximately 20 h. Similar to bioorthogonal labeling of HEK293T cells, surface labeling of ncAA-tagged TARPs was obtained by live incubation with 0.5 μM of cell-impermeable tetrazine-dyes (H-Tet-Cy3, H-Tet-Cy5, Pyr-Tet-ATTO643, Pyr-Tet-AF647). Excess of tetrazine-dye was removed by subsequent washes with warm Tyrode’s solution before imaging of live or fixed neurons (Fig. 1e,f and Fig. 3). Similar to what we observed in HEK293T cells, TCO*A-supplemented neurons transfected with either WT γ2 or WT γ8 displayed no tetrazine-dye labeling (Fig. 3a-c upper panel and d-f upper panel, respectively). Surface labeling of γ2 S44*-positive neurons showed a strong enrichment of γ2 S44* in the dendritic spines with low expression in the dendritic shaft (Fig. 1 g and Fig. 3a-c lower panel). In contrast, γ8 S72* was distributed throughout the dendritic arbor (Fig. 1h and Fig. 3d-f lower panel). Using eGFP as a cell marker, we compared the enrichment of γ2 S44* and γ8 S72* in spines versus dendritic shafts, at the base of the measured spine (extraspine). We observed an average 2.60 ± 0.69 fold higher enrichment in the spines as compared to the neighboring shaft for γ2 S44* (Fig. 3g). Furthermore, γ2 S44* showed a pronounced tendency to accumulate in clusters heterogeneously distributed in the spine head. In contrast, γ8 S72* had a more homogeneous distribution in dendrites and spines with a lower tendency to form clusters. When comparing the fluorescence levels in the spine to the neighboring extraspine area, γ8 S72* was only slightly enriched at the spines (1.17 ± 0.25 fold increase) (Fig. 3g). One of the major drawbacks of overexpression systems, in particular transfection, is the highly heterogeneous expression levels of the protein of interest from cell to cell, which might lead to artifacts like mislocalization of proteins. We thus plotted the mean fluorescence intensity level measured on all analyzed extraspine areas versus the neighboring spine levels. We observed a poor correlation between γ2 S44* labeling at spines versus extraspine, while γ8 S72* displayed a strong correlation between these two areas, independently of the expression level (Fig. 3h). Furthermore, the average enrichment ratio per neuron of both γ2 S44* and γ8 S72* was independent of the expression level (Fig. 3i). This indicates that the difference observed between γ2 S44* and γ8 S72* distribution is independent of their expression level, and likely due to the intrinsic nature of the proteins and their targeting properties.

**Figure 3:**
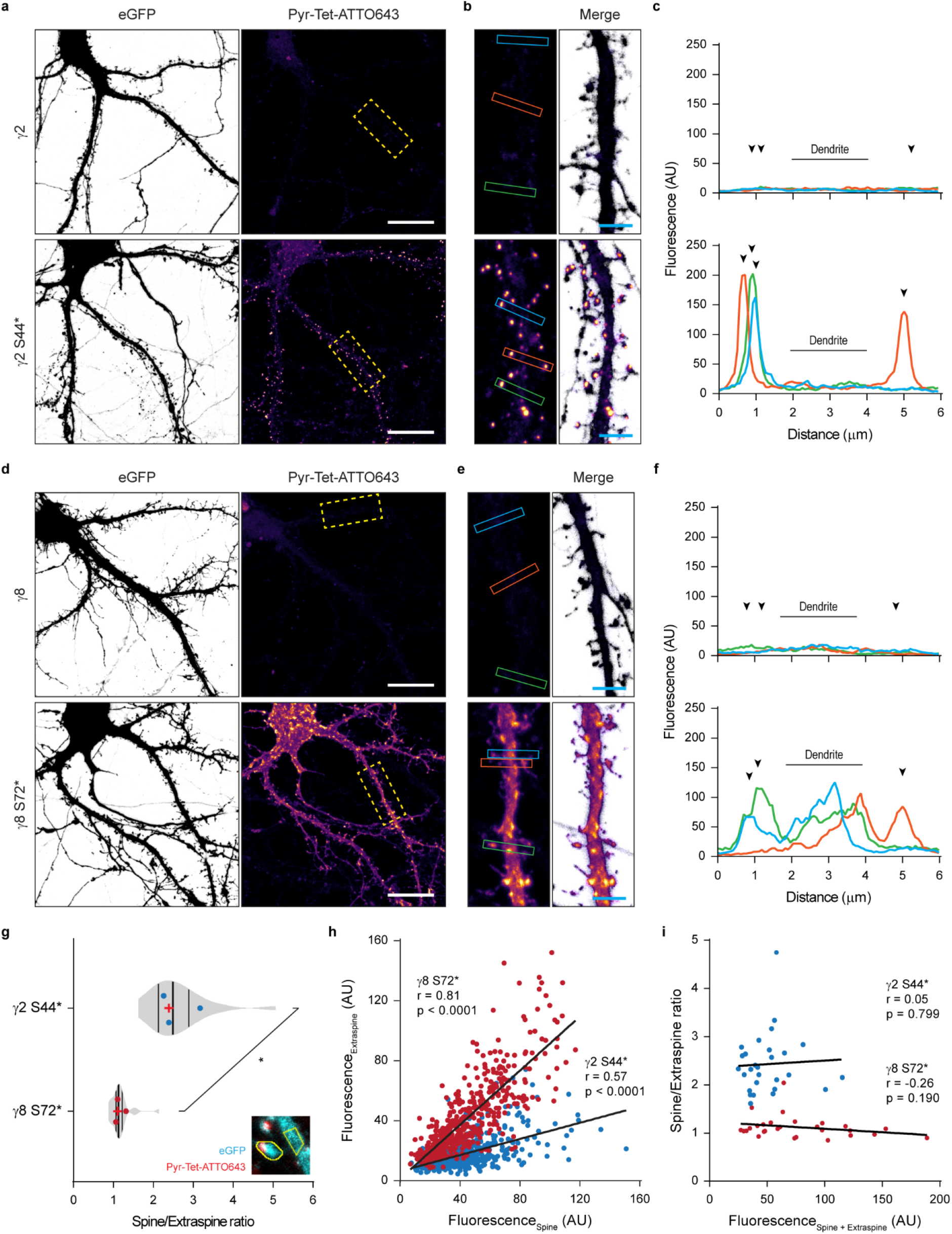
Distinct dendritic surface distribution of γ2 S44* and γ8 S72* in dissociated neurons. (**a-d**) Representative confocal images of fixed dissociated hippocampal neurons coexpressing eGFP, Tet3G/tRNA^Pyl^ and (**a**) pTRE3G-BI PylRS/γ2 (upper panel), pTRE3G-BI PylRS/γ2 S44* (lower panel), (**d**) pTRE3G-BI PylRS/γ8 (upper panel), or pTRE3G-BI PylRS/γ8 S72* (lower panel) in the presence of 250 μM TCO*A and 100 ng.mL^-1^ doxycycline live stained with 0.5 μM Pyr-Tet-ATTO643. (**b**) and (**e**) Magnified views of segments from the respective overview images (dashed rectangles). Scale bar: 20 μm (overview images) and 4 μm (magnified images). (**c**) and (**f**) Line scan measurements of Pyr-Tet-ATTO643 across spines and dendritic shaft based on eGFP signal represented in (**b**) and (**e**). (**g**) Average spine to extraspine intensity ratio of the ncAA staining indicating a spine enrichment of 2.60 ± 0.69 folds for γ2 S44* (blue), and of only 1.17 ± 0.25 fold for γ8 S72* (red). (**h**) Plot of all the analyzed spines fluorescent intensities as a function of the intensity in a corresponding neighboring equivalent extraspine area in the dendrite for γ2 S44* (blue) and γ8 S72* (red) expressing neurons. The Pearson’s correlation coefficients are: γ2 S44*: blue, n = 872, r = 0.57, p < 0.0001; γ8 S72*: red, n = 521, r = 0.81, p < 0.0001. (**i**) Plot of the average ratio per neuron of spine to extraspine intensities as a function of the sum of spine and extraspine intensities for γ2 S44* (blue) and γ8 S72* (red) expressing neurons (~20 spines per neuron).

### Bioorthogonal labeling of TARPs in organotypic hippocampal slice cultures

While dissociated primary neuronal cultures are a well-established experimental model, they lack the physiological cellular environment, network and regional specificity of the intact brain. Given the small size of tetrazine-dyes, high specificity, and ultrafast bioorthogonal reaction with TCO*, we aimed to exploit the potential of this approach as a tool to label surface proteins in the more physiological system of organotypic hippocampal slice cultures (OHSC). We used single-cell electroporation (SCE) to deliver the cDNAs in identified target CA1 pyramidal neurons from 300 μm thick slices. Similar to what we achieved in dissociated neurons, we used the doxycycline-inducible expression system for the controlled expression of ncAA-tagged TARPs and PylRS. TCO*A and doxycycline were added to the media approximately 22 h before tetrazine-dye labeling. Excess of TCO*A and tetrazine-dye in the extracellular space were removed by subsequent washes with warm Tyrode’s solution (Fig. 4a, see Methods section). Confocal images of fixed slices of SCE CA1 neurons co-expressing eGFP and γ2 S44* or γ8 S72* demonstrated good tissue penetrability and high specificity of H-Tet-Cy5 towards TCO* for tissue applications as indicated by the eGFP signal (Fig. 4b,c,e). Similar to what we observed in dissociated neurons (Fig. 3a-f lower panel), γ2 S44* expressed into CA1 neurons showed a remarkable fluorescence signal enrichment at spines of both apical and basal dendrites with reduced labeling in the dendritic shaft (Fig. 4d,g,h). To verify γ2 S44* accumulation along the Z-projected dendritic shaft (Fig. 4d), we co-expressed γ2 S44* with the PSD-95 marker XPH20 fused with eGFP (XPH20::eGFP)^48^ as a reporter and found that γ2 S44* accumulation was indeed always colocalized with the XPH20 eGFP signal (Fig. 4k). In contrary to γ2 S44*, but in line with the observations made in dissociated neurons overexpressing γ8 S72* (Fig. 3d-f lower panel), γ8 S72*-overexpressing CA1 neurons showed a more homogeneously distributed H-Tet-Cy5 fluorescence signal along the dendrites (Fig. 4f,i,j).

**Figure 4:**
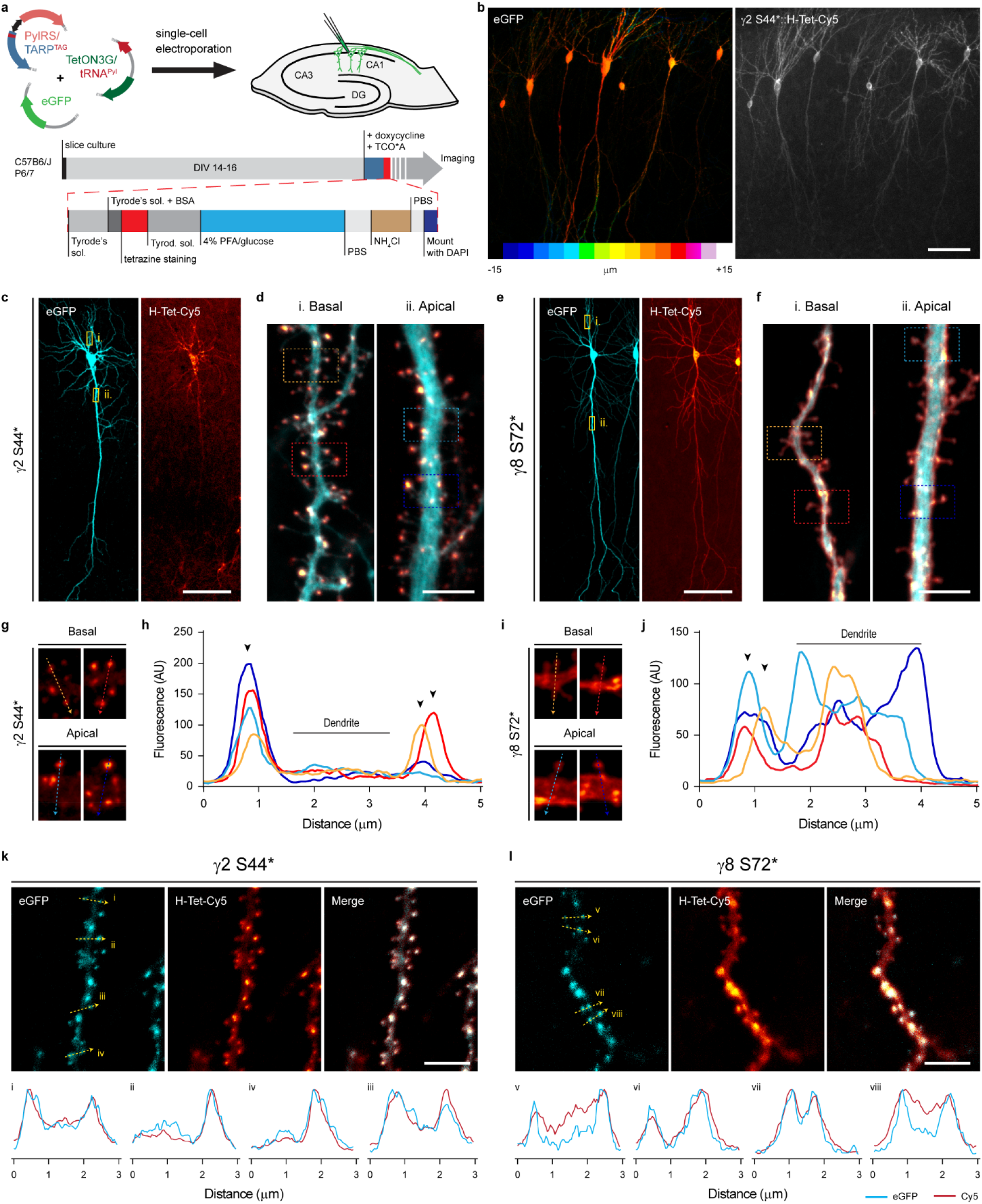
Bioorthogonal labeling of γ2 S44* and γ8 S72* in organotypic hippocampal slice cultures report a distinct surface distribution of TARPs. (**a**) Depiction of the workflow used for expression of ncAA-tagged TARPs in CA1 pyramidal cells in OHSC using single-cell electroporation (SCE), and live staining with tetrazine-dyes. (**b**) Example confocal image of fixed CA1 neurons co-expressing eGFP and γ2 S44* in OHSC. Images are projections of a z-stack taken by 1 μm increments, eGFP signal is color-coded with respect to sample depth. (**c, e**) Representative confocal images of CA1 neurons co-expressing eGFP, and (**c**) γ2 S44*- or (**h**) γ8 S72* live stained with 1 μM H-Tet-Cy5. (**d, f**) Magnified views of segments of the basal and apical dendrites from (**d**) γ2 S44*- and (**f**) γ8 S72*-overexpressing CA1 neurons highlighted in the corresponding overview images (yellow boxes). (**g, i**) Close up of representative spines from (**g**) γ2 S44*- and (**i**) γ8 S72*-overexpressing CA1 neurons highlighted (dashed squares) in the overview images (**d**) and (**f**), respectively. (**h**) and (**j**) Line scan measurements of Cy5 signal across spines in (**g**) and (**i**) respectively. (**k, l**) Confocal images of segments of basal dendrites from CA1 neurons co-expressing either (**k**) γ2 S44* or (**l**) γ8 S72, and the PSD-95 marker, XPH20::eGFP. Bottom insets: line scans of the GFP and Cy5 signal for the 3 μm segments indicated in the above images. Scale bar: 100 μm (overview images) and 5 μm (dendritic segments).

This highlights the reliability of bioorthogonal labeling as a versatile, fast, and specific tool for live labeling of proteins in neuronal tissue.

### *d*STORM imaging reveals differences in nanoscale distribution of TARPs

To investigate the peculiar difference found in the distribution of γ2 S44* and γ8 S72* in neurons by confocal microscopy in more detail, we used SMLM by *direct* stochastic optical reconstruction microscopy (*d*STORM)^49,50^. *d*STORM images revealed the molecular distribution of γ2 S44* and γ8 S72* in hippocampal neurons (Figs. 5a,b) and indicated that γ2 S44* accumulates in synaptic spines (Figs. 5b-d), in agreement with confocal data. To quantify the distribution of ncAA-tagged and H-Tet-Cy5 clicked TARPs, we co-expressed again the PSD-95 marker XPH20::eGFP as a reporter to identify synaptic sites and compared the localization densities determined from *d*STORM data of extrasynaptic and synaptic sites. While both TARPs show a homogeneous distribution in extrasynaptic sites, the absolute localization density determined for γ8 S72* is ~3-fold higher (Fig. 5c). Together with the slightly higher localization density of γ2 S44* in synaptic sites (Fig. 5c), our data thus demonstrate that the localization density measured for y2 S44* is ~9 fold higher in synaptic as compared to extrasynaptic sites, whereas γ8 S72* exhibits only a ~2 fold higher localization density in synaptic compared to extrasynaptic sites (Fig. 5 c, inset).

**Figure 5:**
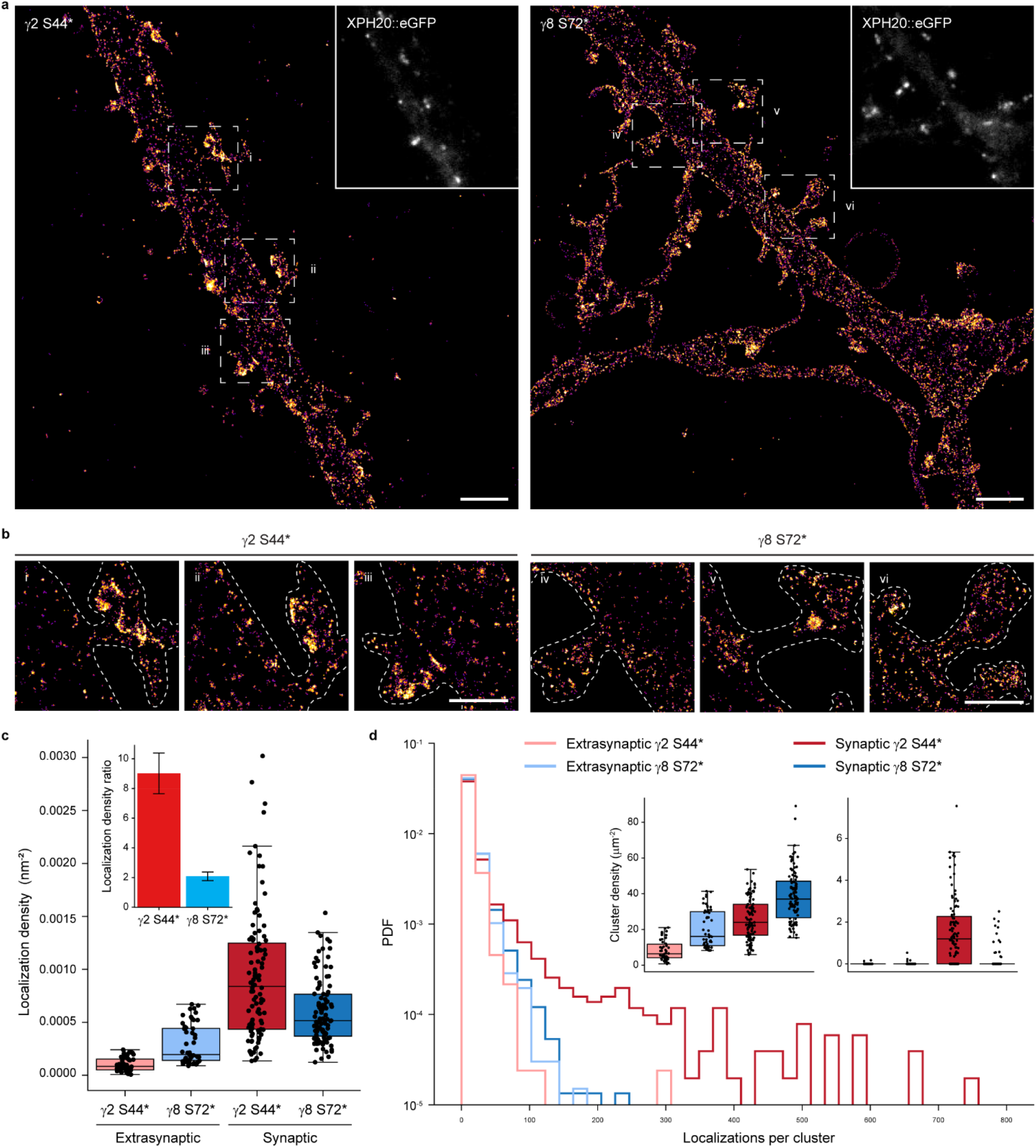
*d*STORM imaging and analysis reveal nano-scale organization of bioorthogonal labeled γ2 S44* and γ8 S72* in dissociated neurons. (**a**) Representative *d*STORM image of Pyr-Tet-AF647 (0.5 μM) live labeled neurons expressing γ2 S44* or γ8 S72* co-expressed with Xph20::eGFP. Scale bar: 2 μm. (**b**) Magnified views of spine and dendrite of the respective overview images in (**a**) (dashed rectangles). Scale bar: 1 μm. (**c**) Boxplots displaying slightly higher synaptic localization densities for γ2 S44* ((0.93 ± 0.06)*10E-3 nm^-2^, n = 104, dark red) compared to γ8 S72* ((0.60 ± 0.03)*10E-3 nm^-2^, n = 102, dark blue). γ8 S72* showed higher extrasynaptic localization densities ((0.29 ± 0.03)*10E-3 nm^-2^, n = 52, light blue) in comparison to γ2 S44* ((0.10 ± 0.01)*10E-3 nm^-2^, n = 50, light red). Inset: Ratio of synaptic to extrasynaptic localization densities indicate a spine enrichment of 9.0 ± 1.4 folds for γ2 S44* (red) and of only 2.1 ± 0.3 for γ8 S72* (blue). (**d**) Histograms showing the number of localizations per cluster for synaptic (dark) and extrasynaptic (light) γ2 S44* (red) and γ8 S72* (blue), displayed as probability density function (PDF). Insets display boxplots of the ROI cluster densities for a selection of clusters with less (left inset) and more than 100 clustered localizations (right inset). Only synaptic γ2 S44* shows clusters with >100 localizations (1.93 ± 0.19 μm^-2^) compared to nearly no clusters for synaptic γ8 S72* (0.21 ± 0.06 μm^-2^) and extrasynaptic γ2 S44* (0.01 ± 0.01 μm^-2^) or γ8 S72* (0.03 ± 0.01 μm^-2^). For a selection of clusters with <100 localizations, γ8 S72* presents larger densities in synaptic (38 ± 1 μm^-2^) as well as extrasynaptic areas (20 ± 2 μm^-2^) in comparison to γ2 S44* clusters (synaptic: 26 ± 1 μm^-2^, extrasynaptic: 8 ± 0.8 μm^-2^). All data represent mean ± standard error of the mean.

Next, we calculated Ripley’s K-function of several regions of interest (ROIs) in synaptic and extrasynaptic areas to analyze the distribution of TARPs in neurons (Supplementary Fig. 3) and compared them to simulated data with spatial distributions following complete spatial randomness or a clustered Neyman-Scott process (accounting for multiple localizations from each fluorophore) in identical ROIs. Ripley’s K-functions showed for both TARPs in and outside of synapses randomly distributed localization clusters with a size of ~20 nm, which can be attributed to multiple localized Cy5 dye molecules. Only γ2 S44* in synaptic areas showed strong deviation from the simulations with a maximum at ~100 nm indicating cluster formation (Supplementary Fig. 3). Individual cluster analysis for each ROI in and outside of synapses confirmed the existence of γ2 S44* cluster in synapses and the absence of extrasynaptic γ2 S44* and γ8 S72* cluster (Fig. 5d). The synaptic ROIs exhibit a higher localization density for γ2 S44* and cluster with average size of ~80 nm (Figs. 5c,d and Supplementary Fig. 4a).

## DISCUSSION

The ability to label target proteins with small ligands, and at sterically hard-to-access epitopes, represents an important challenge in biology, in particular for live-cell and super-resolution imaging studies in neurons. TARPs represent an interesting case study as their limited extracellular loops and close association with AMPARs has prevented the development of adequate ligands, in particular for the study of TARPs organization and trafficking at the cell surface of living neurons. Our motivation to search for alternative labeling strategies was further reinforced by our initial finding that functional antibodies to the extracellular domains of γ2 and γ8 TARPs were unable to recognize native TARPs in neurons, likely due to epitope masking. We thus engineered the technology to incorporate ncAAs in these proteins at given edited sites by GCE to label them directly with fluorophores by click chemistry. We demonstrate that γ2 and γ8 can be directly labeled with this approach both in dissociated primary cultures and organotypic slices of rodent hippocampal neurons. Labeled proteins can then be imaged by widefield, confocal or *d*STORM super-resolution imaging. Our data reveal that γ2 and γ8 display profoundly different distributions on the neuronal surface, γ2 being much more concentrated and clustered at synapses than γ8. In addition, the ability to label the masked epitopes in close proximity to the associated AMPAR subunits allowed us to develop FRET pairs between γ2 or γ8 and GluA subunits.

In the attempt to label surface TARPs, we first developed antibodies against the extracellular domains of γ2 and γ8. While live labeling with our Abs against γ2 Ex2 or γ8 Ex1 was able to specifically detect respectively recombinant γ2 or γ8 in cell lines and dissociated neurons, this approach failed to detect endogenous TARPs in dissociated hippocampal neurons or recombinant γ2 genetically tethered to GluA2. This indicates that the extracellular loops of γ2 and γ8 are masked when associated or in contact with AMPAR, which is compatible with the published cryo-EM structures of TARP/GluA subunit complexes^33–35^. It further indicates that, at endogenous levels, most if not all γ2 and γ8 are associated with AMPAR on the surface of hippocampal neurons, as γ2 in particular only associates to AMPAR subunits^38^. This had remained an important open question in the field. Small tags like hemagglutinin (HA, 9 aa)^38^ or biotin-acceptor peptide (bAP, 15 aa) have been previously successfully inserted in the Ex1 loop of γ2, while bigger proteins such as mCherry led to an intracellular retention of γ2 possibly due to protein misfolding^8^. Surface labeling of γ2-bAP with streptavidin^8^ or γ2-HA with specific Ab^24,51^ in the extracellular loops had previously been achieved in neurons, but most likely only revealed overexpressed TARP not associated with AMPARs. In contrast, GCE combined with click chemistry labeling allowed the site-specific incorporation of ncAAs that can be functionalized and labeled with small tetrazine-dyes with a size of ~1 nm^16^. Both patch-clamp and FRET experiments demonstrate that GCE-labeled TARPs are fully functional and can interact normally with GluA subunits. Indeed, electrophysiological recordings show that the incorporation of ncAAs into the Ex1 of γ2 and γ8 did not compromise TARP-specific AMPAR gating modulation, while FRET experiments indicate close association between ncAA-tagged TARPs and GluA subunits. A parallel approach using cysteine tagging of AMPAR and ncAA-tagging of γ2 TARPs enabled luminescence resonance energy transfer and single-molecule FRET live cell measurements of the distance between GluA2 and γ2 in HEK293T cells and the study of its regulation^32^. Worth mentioning, while we did not observe a difference in terms of labeling efficacy among tested Pyr-Tet-dyes and H-Tet-dyes in both cell lines and dissociated neurons, we did observe that H-Tet-Cy5 outperformed Pyr-Tet-ATTO643 in OHSC (data not shown). This observation can be explained by the faster reaction of H-Tet with TCO*A compared to Pyr-Tet^52,53^. We also observed some decrease in H-Tet-Cy5 fluorescence intensity with depth in organotypic slices, usually accompanied by a decrease in eGFP fluorescence intensity, suggesting inefficient excitation due to scattering issues rather than inefficient tetrazine-dye labeling. Previous electron microscopy studies have suggested γ2 plasma membrane distribution to be almost exclusively synaptic, with γ8 being more equally distributed between extrasynaptic and synaptic sites^39,31^. In addition, functional studies have indicated that γ2 promotes synaptic targeting of AMPARs^19, 29, 24^ whereas γ8 controls extrasynaptic surface pool and synaptic delivery of AMPARs^21,30^. Furthermore, at Schaffer collateral/commissural (SCC) synapses in the adult mouse hippocampal CA1, synaptic inclusion of γ2 potently increases AMPAR expression and transforms low-density synapses into high-density ones, whereas γ8 is essential for low-density or basal expression of AMPARs at non-perforated synapses^39^, which is fully compatible with our observations. Therefore, these TARPs are critically involved in AMPAR density control at SCC synapses. However, specific imaging of γ2 and γ8 distribution in live neurons was lacking due to the absence of adequate tools. Our data indicate that both in dissociated hippocampal and organotypic CA1 pyramidal neurons, γ2 S44* shows a strong accumulation and forms clusters with a size of ~80 nm at spines compared to lower appearance and more homogeneous distribution at the dendritic shaft, while γ8 S72* shows a more homogenous distribution between spines and dendritic shaft without any indication of cluster formation (Fig. 5 c, d). As mentioned, a limitation in our study is the fact that we had to use an overexpression approach. Recent advances in genome editing tools, such as CRISPR/Cas9, will likely make it possible in the future to deliver site-specific incorporation of ncAAs into endogenous proteins in post-mitotic cells, such as neurons. The combination of this approach with future wholegenome recoding in which all the endogenous Amber stop codons are replaced by Ochre codons^54^ would be particularly valuable. Another alternative that might be more reachable in the near future is the use of orthogonal ribosomes^55,56^ combined with quadruplet codons^57^, eliminating the possibility of tRNA-induced suppression of endogenous Amber codons as well as improving the incorporation of ncAA.

In conclusion, the robustness and versatility of the approach shown here, and the panoply of cell-permeable and impermeable tetrazine-dyes^16^ opens a new spectrum of possibilities that remains now to be explored, for multicolor imaging of the nanoscale organization, interactions, and trafficking of intracellular and/or extracellular proteins in living neurons. The minimal perturbation of the target protein by insertion of a ncAA and small size of tetrazine-dyes enables stoichiometric labeling even of sterically shielded protein sites. The method will thus be particularly valuable for quantitative super-resolution microscopy. Altogether, bioorthogonal labeling of TARPs in living neurons constitutes an important achievement in protein tagging in the field of neuroscience, as it not only introduces a robust and fast labeling strategy with minimal to no-perturbation but also allows the labeling of hard-to-tag proteins that to date have been highly affected by the bulky size of previous labeling strategies^58^. This altogether opens the possibility to tackle new sets of biological questions.

## ACKNOWLEDGEMENTS

We thank Edward Lemke and Gemma Estrada Girona (European Molecular Biology Laboratory, Heidelberg, Germany) for the gift of the pCMV tRNA^Pyl^/NESPylRS^AF^ plasmid and expert training on how to use it. We wish to thank AS Hafner and F Coussen for early experiments to develop and characterize the γ2 and γ8 antibodies. We thank the Bordeaux Imaging Center, part of the FranceBioImaging national infrastructure (ANR-10-INBS-04-0) for support in microscopy and in particular C. Poujol for advice and discussions on FLIM-FRET imaging; C. Martin and the IINS in vivo facility for animal husbandry. We thank the IINS cell biology core facilities (LABEX BRAIN [ANR-10-LABX-43]) and in particular C. Breillat and E. Verdier for cell culture and plasmid production. This work was supported by funding from the Ministère de l’Enseignement Supérieur et de la Recherche to D.C., Centre National de la Recherche Scientifique (CNRS), ERC grant ADOS (339541) and DynSynMem (787340) to D.C., grants from the conseil Régional d’Aquitaine to D.C., the MSCA-ITN-ETN SYNDEGEN (675554) to D.C. and D.B.N., Fondation Recherche Médicale (FDT202001010840) to D.B.N.. A.K., G.B. and M.S. acknowledge funding by the Deutsche Forschungsgemeinschaft (DFG, project SA829/19-1) and the European Regional Development Fund (EFRE project “Center for Personalized Molecular Immunotherapy”). S.D. acknowledges funding by the Deutsche Forschungsgemeinschaft (DFG, project DO1257/4-1).

## AUTHOR CONTRIBUTION

D.B.N. performed FRET, patch-clamp and imaging experiments, developed the strategy for GCE in neurons, developed part of the strategies for cDNA constructs and their production, prepared neuronal samples, performed corresponding data analysis and figure preparation and co-wrote the MS., A.K. and G.B. developed strategies for sitespecific ncAA incorporation by GCE, established click mutants, performed mammalian cell culture experiments and performed *d*STORM imaging. A.K., G.B. and S.D. performed the cluster analysis of the *d*STORM data, V.P. co-developed and co-performed the singlecell electroporation in organotypic slices, N.R. developed part of the strategies for cDNA constructs and their production and supervised the neuronal primary culture production, N.C. developed part of the strategies for cDNA constructs and their production, D.P. developed the patch-clamp approach and contributed to the corresponding analysis, M.S. and D.C. co-supervised the study and co-wrote the MS. All authors read and corrected the MS.

## METHODS

### Reagents

Trans-Cyclooct-2-en-L-Lysine (TCO*A; #SC-8008) was purchased from SiChem (Bremen, Germany). Pyrimidyl-Tetrazine-Alexa Fluor 647 (Pyr-Tet-AF647; #CLK-102), Pyr-Tet-ATTO-643 (Pyr-Tet-ATTO643; #CLK-101), H-Tet-Cy3 (#CLK-014-05) and H-Tet-Cy5 (#CLK-015-05) were purchased from Jena Bioscience (Jena, Germany). SNAP-Surface® Alexa Fluor® 488 (BG-AF488; #S9129S) was purchased from New England Biolabs. 2,3-Dioxo-6-nitro-1,2,3,4-tetrahydrobenzo[f]quinoxaline-7-sulfonamide disodium salt (NBQX; #1044) and Kainate (KA; #0222) were purchased from Tocris. L-Glutamic acid monosodium salt (Glu; #G1626) and doxycycline (#D1822) were purchased from Sigma.

### Plasmid constructs

Plasmid amplification was performed via transformation in *E. coli* DH5α (Thermo Fisher Scientific, #EC0111) or *E. cloni®* 10G (Lucigen, #60107) in the case of pTRE3G plasmids, and DNA isolation via MAXI-prep ZymoPURE II Plasmid kits (Zymo Research).

eGFP, mCherry or mEos2 were cloned into the coding sequence of γ2 (between residues 304 and 305) and γ8 (between residues 401 and 402) by introducing AgeI/NheI sites to the respective position. The respective Amber stop mutants (Supplementary Fig. 1c) were generated by introducing a TAG codon through PCR-based site-directed mutagenesis in pcDNA3 vector. For γ8, the endogenous TAG stop codon of WT γ8 was replaced by a TAA stop codon. The plasmid for the expression of the tRNA/aminoacyl transferase pair (pCMV tRNA^Pyl^/NESPylRS^AF^, herein termed PylRS/tRNA^Pyl^) was kindly provided by Edward Lemke^59^.

To reduce TARPs expression toxicity in neurons, and reduce the number of plasmids to transfect, WT TARPs and ncAA-tagged TARPs were subcloned into a bidirectional doxycycline-inducible expression vector pTRE3G-BI (Takara Bio, #631332) using the restriction sites KpnI/XbaI, and the NESPylRS^AF^ was inserted using the restriction sites BamHI/BglII into the BamHI restriction site of the multiple cloning site of pTRE3G-BI after PCR amplification using the oligonocleotides: PylRS_F, 5’-CTTGGATCCGCCACCATGGATAAAAAACC-3’ and PylRS_R, 5’-TAGAAGCTTTTACAGGTTAGTAGAAATACCATTGTAATAG-3’. The U6 promoter and tRNA^Pyl^ were inserted into the pEF1α-Tet3G (Takara Bio, #631336; Tet3G/tRNA) using the restriction site BsrGI after PCR amplification using the oligonucleotides: U6/tRNA_F, 5’-GCATGTACATTTCCCCGAAAAATGG-3’ and U6/tRNA_R, 5’-GGTCATATTGGACATGAGCC-3’ (primer located upstream the U6 promoter on the pCMV tRNAPyl/NESPylRSAF), and coexpressed with the pTRE3G-BI constructs.

γ2 mCherry and tethered GluA2 (flop isoform)::γ2^60^ were subcloned into the doxycycline-inducible expression vector pBI-Tet (Clontech, #6152-1) using the restriction sites MluI/XbaI and MluI/NheI, respectively. pBI-Tet constructs were co-expressed with the pTet-On Advanced Vector (Clontech, #630930).

MfeI/NheI restriction sites were introduced after the signal peptide of GluA1 to insert the SNAPtag® at N-terminus of GluA1 (SNAP::GluA1) flip variant coding sequence in pRK5 vector. The plasmid for the GluA1 Tn5 ME SEP +396 aa was kindly provided by Andrew Plested. AgeI/NheI restriction sites were introduced between the Tn5 ME sequences and SEP was replaced by SNAP-tag® (GluA1::SNAP396). The tethered GluA1::SNAP396::γ2 S61* and GluA1::SNAP396::γ8 K102* were performed as described for the tethered WT GluA1::γ2 in^41^. The plasmid for the expression of the tRNA/aminoacyl transferase pair (pNEU-hMbPylRS-4xU6M15, herein termed PylRS/4xtRNA^Pyl^) was a gift from Irene Coin (Addgene, #105830)^44^. The plasmid for the expression of the Xph20 eGFP CCR5TC (XPH20::eGFP) was a gift from Matthieu Sainlos^48^.

### Heterologous cell culture

HEK293T cells (ECACC, #12022001) were cultured at 37°C under 5% CO_2_ in DMEM supplemented with 10% FBS, 1% L-glutamine and 1% penicillin/streptomycin. COS-7 cells (ECACC, #87021302) were cultured at 37 °C under 5% CO_2_ in DMEM supplemented with 10% FBS, 1% L-glutamine and 1% penicillin/streptomycin.

### Primary dissociated hippocampal neurons

Gestant rat females were purchased weekly (Janvier Labs, Saint-Berthevin, France). Animals were handled and euthanized according to European ethical rules and protocols approved by the local ethics committee office 50. Dissociated hippocampal neurons from embryonic day 18 (E18) Sprague-Dawley rats embryos of either sex were prepared as previously described^61^. Briefly, dissociated neurons were plated at a density of 250,000 cells per 60 mm dish on 0.1 mg.mL^-1^ PLL pre-coated 1.5H, ø 18 mm coverslips (Marienfeld Superior, #0117580). Neurons cultures were maintained in Neurobasal™ Plus Medium (Thermo Fisher Scientific) supplemented with 0.5 mM GlutaMAX (Thermo Fisher Scientific) and 1X B-27™ Plus Supplement (Thermo Fisher Scientific). 2 μM Cytosine β-D-arabinofuranoside (Sigma Aldrich) was added after 72 h. At DIV3/4, cells were transfected with the respective cDNAs using Lipofectamine 2000 (Thermo Fisher Scientific, #11668019). Cultures were kept at 37 °C under 5% CO_2_ up to 18 days.

Astrocytes feeder layers were prepared from the similar embryos, plated between 20,000 to 40,000 cells per 60 mm dish and cultured in Minimum Essential Medium (Thermo Fisher Scientific) containing 4.5 g.L^-1^ glucose, 2 mM GlutaMAX and 10% heat-inactivated horse serum for 14 days.

### Organotypic hippocampal slice cultures (OHSC)

Animals raised in our animal facility were handled and euthanized according to European ethical rules and ehtics committee. OHSC from animals at postnatal day 5-7 from wild type mice of either sex (C57Bl6/J strain) were prepared as previously described^62^. Briefly, animals were quickly decapitated and hippocampi were dissected out and placed in ice-cold carbonated dissection buffer (in mM): 230 sucrose, 4 KCl, 5 MgCl_2_, 1 CaCl_2_, 26 NaHCO_3_, 10 D-glucose, and phenol red. Coronal slices (300 μm) were cut using a tissue chopper (McIlwain), collected and positioned on interface-style Millicell® culture inserts (Millipore) in 6 well culture plates containing 1 mL of sterile serum-containing MEM medium (in mM): 30 HEPES, 5 NaHCO_3_, 0.511 sodium L-ascorbate, 13 D-glucose, 1 CaCl_2_, 2 MgSO_4_, 5 L-glutamine, and 0.033% (v/v) insulin, pH 7.3, osmolarity adjusted to 317-320 mOsm, plus 20% (v/v) heat-inactivated horse serum. Brain slices were incubated at 35 °C under 5% CO_2_ and the culture medium was changed from the bottom of each well every 2 to 3 days. After 14-15 days in culture, slices were transferred to an artificial cerebrospinal fluid containing (in mM): 130 NaCl, 2.5 KCl, 2.2 CaCl_2_, 1.5 MgCl_2_, 10 D-glucose, and 10 HEPES, pH 7.35, osmolarity adjusted to 300 mOsm. CA1 pyramidal cells were then processed for single-cell electroporation (SCE) using glass micropipettes containing K-gluconate-based intracellular solution (in mM): 135 K-gluconate, 4 NaCl, 2 MgCl_2_, 2 HEPES, 2 Na2ATP, 0.3 NaGTP, 0.06 EGTA, 0.01 CaCl_2_ (pH 7.2-7.3 with KOH, osmolarity adjusted to 290 mOsm) with plasmids encoding Tet3G/tRNA^P^y^l^ and pTRE3G-BI PylRS/γ2 S44* or pTRE3G-BI PylRS/γ8 S72* in equal proportions (26 ng. μl^-1^) along with eGFP (13 ng.μl^-1^) or XPH20 eGFP (13 ng.μl^-1^). Patch pipettes were pulled from 1 mm borosilicate capillaries (Harvard Apparatus) with a vertical puller (Narishige, #PC-100). SCE was performed by applying 4 square pulses of negative voltage (−2.5 V, 25 ms pulse width) at 1 Hz. After SCE, slices were placed back in the incubator for 4-5 days before labeling.

### Electrophysiology

cDNAs for GluA1 (250 ng), PylRS/tRNA^RS^ (375 ng), and WT/ncAA-tagged γ2/γ8 eGFP or soluble eGFP (375 ng) were co-transfected into HEK293T cells (90,000-100,000 cells.cm^-2^ in 12-well plate) using jetPRIME® (Polyplus-transfection, #114-01). 250 μM TCO*A and 40 μM NBQX were added to the cells at the time of the transfection. Cells were trypsinized 1 day after transfection and seeded on PLL-coated coverslips. Cells were transferred to the recording chamber, and brightly fluorescent isolated cells were selected. Whole-cell patch-clamp recordings were performed at room temperature in HEPES-buffered Tyrode’s solution (HBSS) containing (in mM): 138 NaCl, 2 KCl, 2 MgCl_2_, 2 CaCl_2_, 10 D-glucose, and 10 HEPES, pH 7.4, osmolarity adjusted to 317-320 mOsm. Patch pipettes were filled with an internal solution containing (in mM): 120 CsCH_3_SO_3_, 2 NaCl, 2 MgCl_2_, 10 EGTA, 100 HEPES, and 4 Na_2_ATP, pH 7.4, osmolarity 312 mOsm. Pipette resistances for these experiments were typically 3-5 MΩ and cells with a series resistance higher than 15 MΩ were discarded. Glu (10 mM) or KA (0.1 mM) were dissolved in HEPES-buffered solution and applied using a theta pipette driven by a piezoelectric controller (Burleigh, #PZ-150M). Membrane potential was held at −60 mV. Currents were collected using an EPC10 amplifier (HEKA) and filtered at 2.9 kHz and recorded at a sampling frequency of 20 kHz.

### TARPs immunostaining

cDNAs for Stg mEos2 or γ8 mEos2 (500 ng) were transfected into COS-7 cells (14,000-17,000 cells/cm^2^ in 12-well plate) for 24 h using X-tremeGENE HP DNA (Roche, #06366236001). Cells were incubated for 7 min at 37 °C with either 4 μg.mL^-1^ rabbit anti-γ2 Ex2 or 1:50 serum rabbit anti-γ8 Ex1 antibodies prior to fixation. Dissociated hippocampal neurons were co-transfected either with pBI-Tet γ2::mCherry or tethered pBI-Tet eGFP::GluA2::γ2 and pTet-On Advanced Vector in equal proportions (125 ng). Transfected neurons were treated with 200 ng.mL^-1^ doxycycline 18 h prior to use. Neurons were incubated for 7 min at 37 °C with 10 μg.mL^-1^ mouse anti-GluA (Synaptic Systems, #182411) and anti-γ2 Ex2 or anti-γ8 Ex1 antibodies prior to 4% PFA/sucrose fixation. Reactive aldehydes groups were blocked for 10 min with 50 mM NH4Cl. Alternatively, neurons were live incubated with the anti-GluA antibody, and after fixation neurons were permeabilized with 0.2% Triton-X100 for 5 min and incubated with 0.4 μg.mL^-1^ rabbit anti-γ8 (Frontiers Institute, #TARPg8-Rb-Af1000) diluted in 3% BSA in PBS. Cells were incubated with the respective secondary antibodies anti-mouse AF568 and anti-rabbit AF647 (Thermo Fisher Scientific) diluted at 1:1000 in 3% BSA in PBS. Imaging was performed on an up-right widefield fluorescence microscope (Leica Microsystems, Leica DM5000 B) microscope controlled by Metamorph software (Molecular Devices). Fluorescence excitation of eGFP, AF568 and AF647 was done by a LED SOLA Light (Lumencor). Images were acquired using an oil-immersion objective (Leica, HCX PL APO 40x/NA 1.25 OIL) and appropriate filter set. Fluorescent emission was collected using a sCMOS camera (Hamamatsu Photonics, ORCA-Flash4.0 V2).

### Bioorthogonal labeling in HEK293T cells

HEK293T cells plated at a density of 80,000-90,000 cells.cm^-2^ on a pre-coated PDL 4-well Nunc™ Lab-Tek™ II chamber (Thermo Fisher Scientific, #155382PK) were co-transfected with PylRS/4xtRNA^Pyl^ (500 ng) and respective tagged TARPs (500 ng) using jetPRIME^®^ transfection reagent for 24 h in the presence or absence of 250 μM TCO*A. Cells were washed once with cell media to remove excessive TCO*A prior to labeling with 1.5 μM Pyr-Tet-ATTO643 or H-Tet-Cy5 diluted in TCO*A-free medium for 30 min on ice. Subsequently, cells were rinsed 3 times with icecold HBSS and immediately live imaged or fixed for 15 minutes at RT with 4% FA in PBS followed by 3 washing steps with HBSS before imaging. Confocal imaging of living or fixed cells was performed using a LSM700 setup (Zeiss) equipped with an oil-immersion objective (Zeiss, PlanApochromat 63x/NA 1.4 OIL). eGFP and Pyr-Tet-ATTO643/H-Tet-Cy5 were excited using a 488 nm or 641 nm solid-state laser and respective filter settings. Images were processed in ImageJ (FIJI) adjusting brightness and contrast to identical values for comparison of experiments.

### Bioorthogonal labeling in dissociated hippocampal neurons

Dissociated hippocampal neurons were co-transfected with Tet3G/tRNA^Pyl^ (104 ng), pTRE3G-BI PylRS/TARPs (γ2, γ8, γ2 S44* or γ8 S72*; 104 ng), along with eGFP or XPH20 eGFP (42 ng) at DIV 3/4 using lipofectamine 2000. At DIV16-18, 250 μM TCO*A and 100 ng.mL^-1^ doxycycline were added to the medium. After ~20 h, cells were rinsed 3 times with warm Tyrode’s solution containing (in mM): 100 NaCl, 5 KCl, 5 MgCl_2_, 2 CaCl_2_, 15 D-glucose, and 10 HEPES, pH 7.4, osmolarity adjusted to 243-247 mOsm followed by 3 min incubation in Tyrode’s solution containing 1% BSA. Cells were then incubated with 0.5 μM tetrazine-dye for 7 min at 37 °C and rinsed 4 times with Tyrode’s solution.

Live-cell imaging was performed in Tyrode’s solution at 37 °C using an incubator box with an air heater system (Life Imaging Services) installed on an inverted Leica DMI6000 B (Leica Microsystem) spinning disk microscope controlled by Metamorph software (Molecular Devices). Z-stacks of whole neurons were acquired using an oil-immersion objective (Leica, HCX PL APO 40x/NA 1.25 OIL) and appropriate filter set. Fluorescent emission was collected using a sCMOS camera (Hamamatsu, ORCA-Flash4.0 V2).

Alternatively, cells were fixed for 10 min using 4% PFA/glucose. Reactive aldehydes groups were blocked for 10 min with 50 mM NH4Cl. Images of fixed neurons were acquired with a Leica TCS SP8 confocal microscope controlled by Leica Application Suite X (LAS X) software and equipped with hybrid detectors. eGFP and Pyr-Tet-ATTO643 were excited at 488 nm and 638 nm, respectively. For quantification of γ2 S44* and γ8 S72* surface distribution in dissociated hippocampal neurons, Z-stacks of whole dendrite segments were acquired using an oil-immersion objective (Leica, HC PL APO CS2 63x/NA1.40 OIL) and a pinhole opened to one time the Airy disk.

### Bioorthogonal labeling in OHSC

Single electroporated neurons from OHSC co-expressing pTRE3G-BI PylRS/γ2 S44* or pTRE3G-BI PylRS/γ8 S72*, Tet3G/tRNA^Pyl^ and eGFP were treated with 250 μM TCO*A and 100 ng.mL^-1^ doxycycline for ~22 h before labeling. Slices were washed three times 5 min with warm Tyrode’s solution followed by 5 min in Tyrode’s solution containing 1% BSA. Subsequently, slices were incubated for 10 min at 35 °C with 1 μM H-Tet-Cy5 diluted in Tyrode’s solution containing 1% BSA and washed four times 5 min with Tyrode’s solution. Slices were fixed for 2 h at RT with 4% PFA/sucrose, washed with PBS. Reactive aldehydes groups were blocked for 20 min in 200 mM NH4Cl. Slices were mounted in Fluoromount-G Mounting Medium (Thermo Fisher Scientific, #00-4958-02) and left to cure for 48 h at RT before imaging.

Images of fixed neurons were acquired with a Leica TCS SP8 confocal microscope controlled by Leica Application Suite X (LAS X) software and equipped with hybrid detectors. eGFP and Pyr-Tet-ATTO643 were excited at 488 nm and 638 nm, respectively. Z-stacks of whole neuron were acquired using an oil-immersion objective (Leica, 20x/NA 0.70 IMM) and a pinhole opened to two times the Airy disk. For quantification of γ2 S44* and γ8 S72* surface distribution, Z-stacks of segments basal and apical dendrite were acquired using an oil-immersion objective (Leica, HC PL APO CS2 63x/NA 1.40 OIL) and a pinhole opened to one time the Airy disk.

### γ2 S44* and γ8 S72* surface distribution in neurons

All images were analyzed using ImageJ (FIJI) software. Confocal images of dissociated neurons co-expressing eGFP, Tet3G/tRNA^Pyl^ and pTRE3G-BI PylRS/TARPs (γ2, γ8, γ2 S44* or γ8 S72*) were maximum intensity Z-projected. For tetrazine specificity, 3 pixel-width line scans across spines and dendritic shaft and cell-free areas were performed based on eGFP fluorescence. For surface distribution, masks of regions of interest (spine and adjacent dendritic draft area) generated based on thresholded eGFP images upon a Gaussian blur filter (radius = 1) were applied. Spine enrichment was calculated as the mean spine fluorescence intensity over the neighbor dendritic area mean fluorescence.

For surface distribution of γ2 S44*, γ8 S72*, and XPH20::eGFP in OHSC, confocal images of dendritic segments were integrated intensity Z-projected. Upon a Median filter (radius = 1) was applied, 3 pixel-width line scans across spines that were perpendicular to the dendritic shaft were performed based on eGFP fluorescence.

### *d*STORM imaging

The TARP constructs γ2 S44* or γ8 S72*-positive neurons at DIV17-18 co-transfected with XPH20::eGFP were live stained with 0.5 μM Pyr-Tet-AF647 and fixed with 4% FA and 0.25% GA in PBS for 15min.

The dSTORM images were acquired using an inverted wide-field fluorescence microscope (Olympus, IX-71). For excitation of Pyr-Tet-AF647 a 640-nm optically pumped semiconductor laser (OPSL) (Chroma, Genesis MX639-1000 STM, Coherent, Cleanup 640/10) was focused onto the back focal plane of the oil-immersion objective (Olympus, 60x, NA 1.45). Emission light was separated from the illumination light using a dichroic mirror (Semrock, FF 410/504/582/669 Brightline) and spectrally filtered by a bandpass filter (Semrock, 679/41 BrightLine HC). Images were recorded with an EMCCD (Andor, Ixon DU897). Resulting pixel size for data analysis was measured as 129 nm. For each dSTORM measurement, at least 15,000 frames at 50 Hz and irradiation intensities of ~2 kW cm^-2^ were recorded by TIRF (total internal reflection fluorescence) illumination. Experiments were performed in PBS-based photoswitching buffer containing 100 mM β-mercaptoethylamine (MEA; Sigma-Aldrich) adjusted to pH 7.4. Image reconstruction was performed using rapidSTORM3.3^63^. Overview images were reconstructed with pixel size of 20 nm, whereas insets were calculated with 10 nm pixel size. Prior to *d*STORM imaging, fluorescent image of XPH20::eGFP was acquired at 10 Hz using a 487 nm diode laser (TopticaPhotonics, iBEAM-SMART-488-S-HP), a dichroic mirror (Semrock, FF 410/504/582/669 Brightline) and a bandpass filter (Chroma, ET525/50).

### *d*STORM imaging analysis

Cluster analysis was conducted using a custom custom-written python script applying DBSCAN algorithm as well as Ripley K analysis on localization data in determined region of interests (ROIs). In advance, Xph20::eGFP images were merged in ImageJ (Fiji) with the corresponding superresolved reconstructed image to identify synaptic and extrasynaptic areas. Contrast and brightness of eGFP signal was dilated using ImageJ to determine ROIs of similar size in neuronal spines for y2 S44* and y8 S72* (Supplementary Fig. 4 b). Synaptic and extrasynaptic localization densities describe the number of localizations detected per ROI area. All dSTORM analysis was carried out on localizations in frames between 2 000 and 15 000, with intensity of more than 6500 camera counts and with a local background of less than 800. DBSCAN (with parameter epsilon of 20 nm and minPoints of 3) was applied for identification of clustered localizations of TARPs. Distributions for localizations per cluster and cluster area of synaptic and extrasynaptic y2 S44* as well as y8 S72* were displayed by their probability density function. The cluster density (number of clusters per ROI area) was calculated for clusters with less and more than 100 localizations per cluster. Cluster analysis was performed on 5 neurons of y2 S44* (3 independent experiments) and y8 S72* (4 independent experiments) resulting in analysis of synaptic y2 S44* ROIs (n = 104), synaptic y8 S72* ROIs (n = 102), extrasynaptic y2 S44* ROIs (n = 50) and extrasynaptic y8 S72* ROIs (n = 52).

We calculated and displayed Ripley’s H-function, a normalized Ripley’s K-function, as previously described^64, 65^. Computation was carried out for each ROI without edge correction. The averaged H-function was compared to H-functions and their 95% confidence intervals were computed from 100 simulated data sets with localizations distributed on the same ROIs (and identical number of localizations in each ROI) according to complete spatial randomness or a Neyman-Scott process. The Neyman-Scott clustering process has homogeneously distributed parent events with each parent having n offspring events, where n is Poisson distributed with mean 10, and with the offspring positions having a Gaussian offset with a standard deviation of 12 nm. The maximum of the H-function indicates a distance that is between cluster radius and diameter and thus provides an estimate for the average cluster size.

### Frequency domain-based fluorescence lifetime imaging (FLIM)-Förster resonance energy transfer (FRET) measurements

HEK293T cells plated at a density of 50,000-60,000 cells.cm^-2^ on a pre-coated PLL 4-well Nunc™ Lab-Tek™ II chamber were co-transfected with PylRS/tRNA^Pyl^ (166 ng), ncAA-tagged TARPs (166 ng) and SNAP-tagged GluA1 (166 ng), or PylRS/tRNA^Pyl^ and tethered GluA1 SNAP396::γ2 S61* or GluA1 SNAP396::γ8 K102* in equal amounts (250 ng) using jetPRIME®. 250 μM TCO*A and 40 μM NBQX were added to the cells at the time of the transfection. After 48 h, cells were incubated with 1.5 μM H-Tet-Cy3 and 5 μM BG-AF488 diluted in TCO*A-free medium for 30 min at 37°C. Cells were rinsed three times with HBSS.

Experiments were performed in HBSS at 37°C using an incubator box with an air heater system (Life Imaging Services) installed on an inverted Leica DMI6000 B (Leica Microsystem) spinning disk microscope and using the LIFA frequency-domain lifetime attachment (Lambert Instruments) and the LI-FLIM software. Cells were imaged with an oil-immersion objective (Leica, HCX PL Apo 100x/NA 1.4 oil) using an appropriate GFP filter set. Cells were excited using a sinusoidally-modulated 3 W 477 nm light-emitting diode at 40 MHz under widefield illumination. Fluorescence emission was collected using an intensified CCD LI2CAM MD camera (Lambert Instruments, FAICM). Lifetimes were referenced to a 1 mg.mL^-1^ erythrosine B that was set at 0.086 ns^66^. The lifetime of the sample was determined from the fluorescence phase-shift between the sample and the reference from a set of 12 phase settings using the manufacturer’s LI-FLIM software. All data are pulled measurements from a minimum of 20 cells per individual preparation. At least 20 cells in a minimum of three individual preparations were taken in consideration, except GluA1 SNAP396::γ2 S44*-AF488/Cy3 which are from two preparations.

### Statistics

All electrophysiological recordings were analyzed with IGOR Pro 5 (WaveMetrics). Current amplitudes were measured with built-in tools, and τ_des_ was measured with exponential fit using a least-squares algorithm.

Statistical significance was calculated using GraphPad Prism. Statistical values are given as mean ± SD or SEM (as indicated); ***p < 0.001, **p < 0.01, *p < 0.05, n.s. p > 0.05. Box and violin plot represent median, lower and upper quartiles, mean (cross), min to max values (electrophysiology data) or 1.5 times the interquartile range (*d*STORM data) (box plot whiskers), and individual values (electrophysiology and *d*STORM data) or mean of independent experiments (FRET data and Fig. 3g) (dots). For multiple sample comparisons within electrophysiology experiments, one-way ANOVA with a Fisher’s Least Significant Difference multiple comparisons test was used. For multiple sample comparisons within the FRET experiments, one-way ANOVA with a Tukey’s multiple comparisons test was used on the mean AF488 lifetime for each condition from a set of 2 to 5 independent experiments.

### Data and code availability statement

The datasets generated during and/or analyzed during the current study are available from the corresponding author on reasonable request. Custom Python code for analysis procedures can be made available upon reasonable request to the corresponding author.

---

**Supplementary Figure 1:**
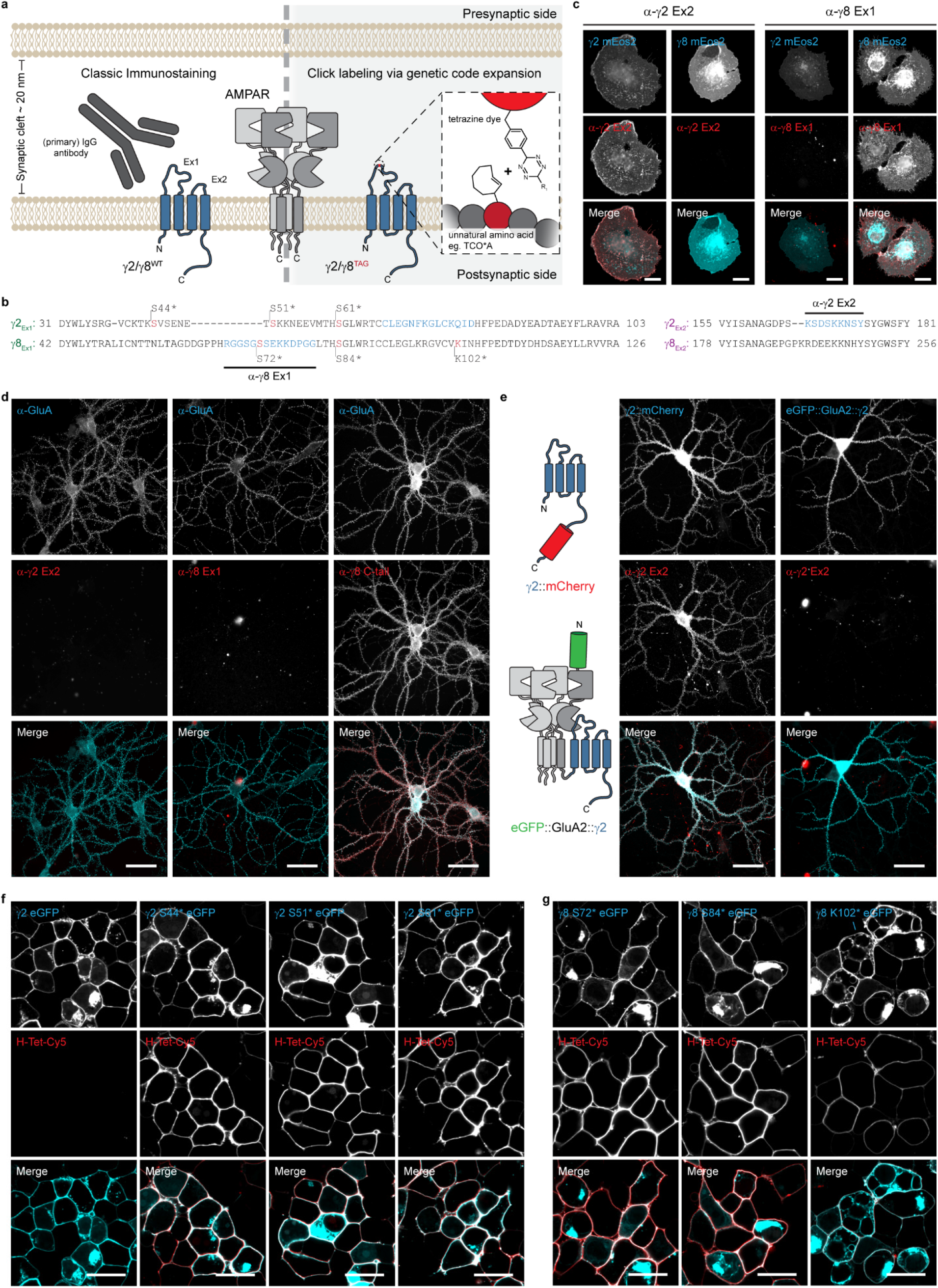
Unmasking the EX1 loop of TARPs using bioorthogonal labeling. (**a**) Schematic illustration of the two approaches used to label the extracellular pool of γ2 and γ8. On the left side, the classic indirect immunostaining using whole IgG antibodies, and on the right, click chemistry labeling via genetic code expansion. (**b**) Sequence alignment of the extracellular loops, Ex1 and Ex2, of γ2 and γ8 from *rattus norvegicus*. Amber substitution mutations are represented in red. The epitopes recognized by the antibodies are represented in blue. (**c**) Representative widefield images of fixed COS7 expressing either γ2 or γ8 bearing mEos2 live stained with the antibodies against the extracellular loops of γ2 (α-γ2 Ex2) or γ8 (α-γ8 Ex1). (**d**) Representative widefield images of fixed untransfected dissociated hippocampal neurons costained live with α-GluA1/2/3/4, and α-γ2 Ex2 (left), α-γ8 Ex1 (middle), or post-fixation/permeabilization with α-γ8 C-tail (right). (**e**) Left: schematic illustration of the two tagged-γ2 used for their expression, respectively γ2 mCherry (upper) and eGFP GluA2::γ2 (lower). Right: representative widefield images of fixed dissociated hippocampal neurons co-expressing pTet-On Advanced Vector, and the doxycyline-inducible pBI-Tet γ2::mCherry (left) or pBI-Tet eGFP GluA2::γ2, live stained with α-γ2 Ex2. (**f-g**) Representative confocal images of live HEK293T cells co-expressing PylRS/4xtRNA^Pyl^, and (**f**) γ2::eGFP or ncAA-tagged γ2::eGFP or (**g**) ncAA-tagged γ8::eGFP in the presence of 250 μM TCO*A stained with 1.5 μM Pyr-Tet-ATTO643. Scale bar: (**b**), (**f**) and (**g**) 20 μm, (**c**) and (**d**) 50 μm.

**Supplementary Figure 2:**
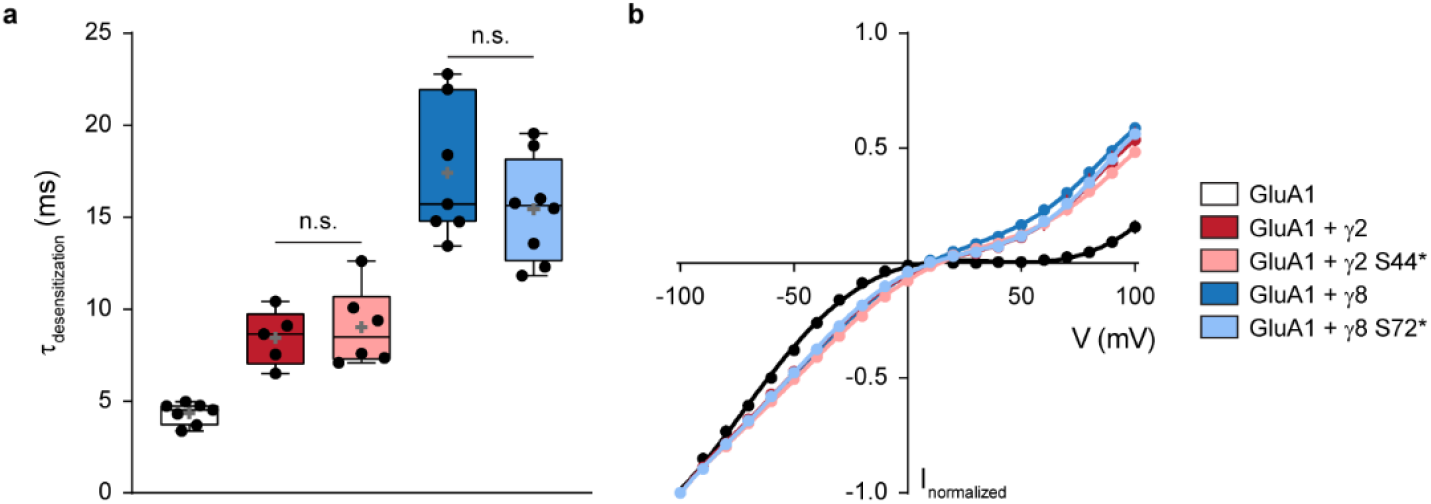
Incorporation of TCO*A within the EX1 does not affect TARP subtype-induced AMPAR modulation. (**a**) Desensitization rates in response to 100 ms of 10mM Glu applied to whole-cell patches from HEK293T cells co-expressing PylRS/tRNA^Pyl^, GluA1 and, eGFP (τ_des_ = 4.34 ± 0.06 ms; n = 7; black), γ2::eGFP (τ_des_ = 8.44 ± 1.50 ms; n = 5; blue), γ2 S44*::eGFP (τ_des_ = 9.02 ± 2.14 ms; n = 6; light blue), γ8::eGFP (τ_des_ = 17.41 ± 3.72 ms; n = 7; red), and γ8 S72*::eGFP (τ_des_ = 15.43 ± 2.82 ms; n = 8; light red). All data represent mean ± standard deviation. (**b**) I-V relationships for 10 mM Glu-evoked peak currents applied to whole-cell patches from HEK293T cells co-expressing PylRS/tRNA^Pyl^, GluA1 and, eGFP (control; n = 7; black), γ2::eGFP (n = 5; blue), γ2 S44*::eGFP (n = 6; light blue), γ8::eGFP (n = 7; red), and γ8 S72*::eGFP (n = 8; light red). Current are normalized to −100 mV.

**Supplementary Figure 3:**
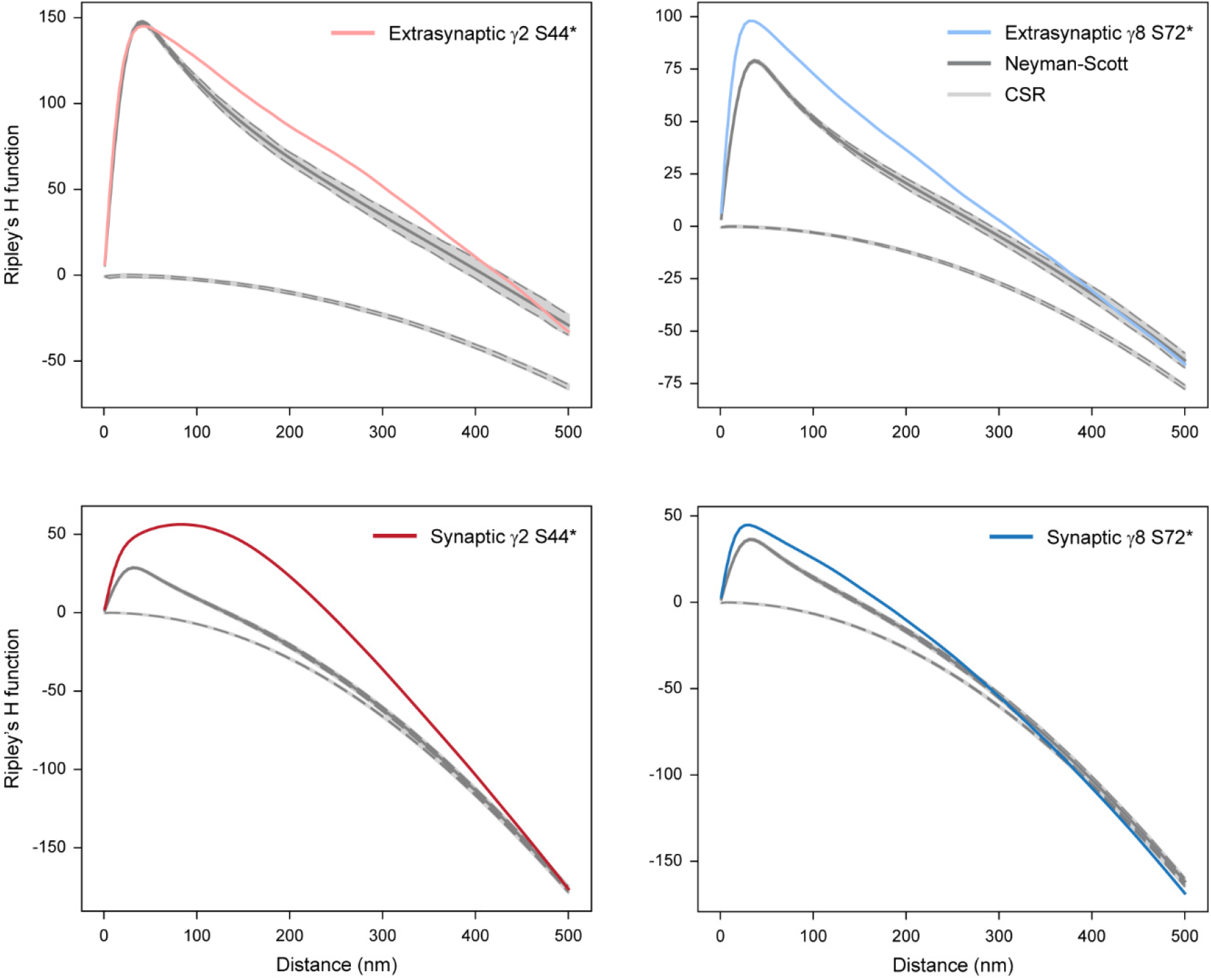
Ripley’s H function. γ2 S44* (red) as well as γ8 S72* (blue) show synaptic (dark) and extrasynaptic (light) random distributions of localization clusters with a size of ~20 nm. Only γ2 S44* (dark red) shows a non-random distribution in synaptic areas with a maximum at ~100 nm indicating cluster formation. Ripley’s H function from 100 replicates of simulated data with spatial distributions following complete spatial randomness (lower grey lines) or a clustered Neyman-Scott process (upper grey lines) in identical ROIs are displayed with 95% confidence intervals (dotted gray lines).

**Supplementary Figure 4:**
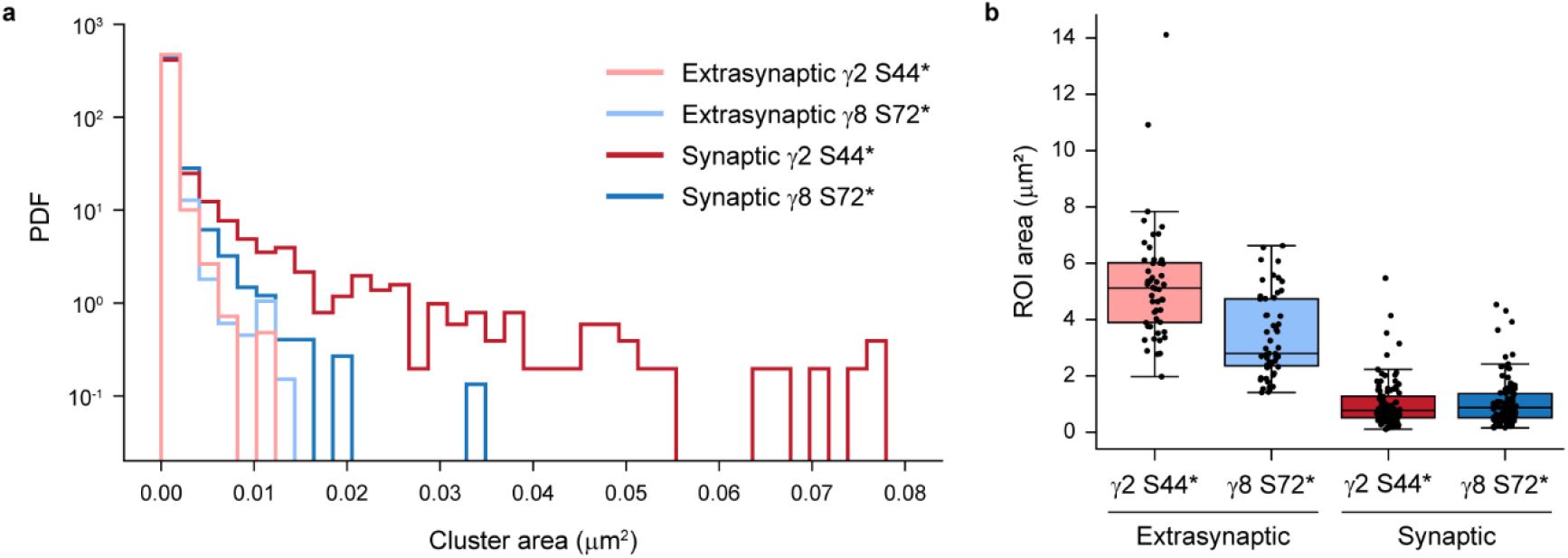
DBSCAN analysis of γ2 S44* and γ8 S72* localizations reveal the presence of clusters (>80nm) only for γ2 S44*in synapses. (**a**) Histograms of cluster areas displaying the probability density function (PDF). Extrasynaptic γ8 S72* (light blue), extrasynaptic γ2 S44* (light red) and synaptic γ8 S72* (dark blue) show similar cluster areas for all clusters with areas < 0.02 μm^2^, whereas synaptic γ2 S44* (dark red) exhibit also clusters with cluster areas > 0.02 μm^2^ corresponding to cluster diameter > ~80 nm. (**b**) Boxplots displaying selected ROI areas that were included in cluster analysis. ROI areas were expanded to be similar for synaptic γ8 S72* and γ2 S44* as well as for extrasynaptic γ8 S72* and γ2 S44* to ensure comparable cluster analysis between γ8 and γ2.

## REFERENCES

1. Hussey, A.M. & Chambers, J.J. Methods to locate and track ion channels and receptors expressed in live neurons. ACS Chem Neurosci 6, 189–198 (2015).

2. Groc, L. & Choquet, D. Linking glutamate receptor movements and synapse function. Science 368 (2020).

3. Sigal, Y.M., Zhou, R. & Zhuang, X. Visualizing and discovering cellular structures with superresolution microscopy. Science 361, 880–887 (2018).

4. Sauer, M. Localization microscopy coming of age: from concepts to biological impact. J Cell Sci 126, 3505–3513 (2013).

5. Sauer, M. & Heilemann, M. Single-Molecule Localization Microscopy in Eukaryotes. Chem Rev 117, 7478–7509 (2017).

6. Weber, K., Rathke, P.C. & Osborn, M. Cytoplasmic microtubular images in glutaraldehyde-fixed tissue culture cells by electron microscopy and by immunofluorescence microscopy. Proc Natl Acad Sci U S A 75, 1820–1824 (1978).

7. Zwettler, F.U. et al. Molecular resolution imaging by post-labeling expansion single-molecule localization microscopy (Ex-SMLM). Nat Commun 11, 3388 (2020).

8. Chamma, I. et al. Mapping the dynamics and nanoscale organization of synaptic adhesion proteins using monomeric streptavidin. Nat Commun 7, 10773 (2016).

9. Virant, D. et al. A peptide tag-specific nanobody enables high-quality labeling for dSTORM imaging. Nat Commun 9, 930 (2018).

10. Rothbauer, U. et al. Targeting and tracing antigens in live cells with fluorescent nanobodies. Nat Methods 3, 887–889 (2006).

11. Opazo, F. et al. Aptamers as potential tools for super-resolution microscopy. Nat Methods 9, 938–939 (2012).

12. Schlichthaerle, T. et al. Site-Specific Labeling of Affimers for DNA-PAINT Microscopy. Angew Chem Int Ed Engl 57, 11060–11063 (2018).

13. Liu, C.C. & Schultz, P.G. Adding new chemistries to the genetic code. Annu Rev Biochem 79, 413–444 (2010).

14. Chin, J.W. Expanding and reprogramming the genetic code of cells and animals. Annu Rev Biochem 83, 379–408 (2014).

15. Nikic, I., Kang, J.H., Girona, G.E., Aramburu, I.V. & Lemke, E.A. Labeling proteins on live mammalian cells using click chemistry. Nat Protoc 10, 780–791 (2015).

16. Beliu, G. et al. Bioorthogonal labeling with tetrazine-dyes for super-resolution microscopy. Commun Biol 2, 261 (2019).

17. Ernst, R.J. et al. Genetic code expansion in the mouse brain. Nat Chem Biol 12, 776–778 (2016).

18. Osten, P. & Stern-Bach, Y. Learning from stargazin: the mouse, the phenotype and the unexpected. Curr Opin Neurobiol 16, 275–280 (2006).

19. Schnell, E. et al. Direct interactions between PSD-95 and stargazin control synaptic AMPA receptor number. Proc Natl Acad Sci U S A 99, 13902–13907. (2002).

20. Choquet, D. & Hosy, E. AMPA receptor nanoscale dynamic organization and synaptic plasticities. Curr Opin Neurobiol 63, 137–145 (2020).

21. Rouach, N. et al. TARP gamma-8 controls hippocampal AMPA receptor number, distribution and synaptic plasticity. Nat Neurosci 8, 1525–1533 (2005).

22. Park, J. et al. CaMKII Phosphorylation of TARPgamma-8 Is a Mediator of LTP and Learning and Memory. Neuron 92, 75–83 (2016).

23. Chen, L. et al. Stargazin regulates synaptic targeting of AMPA receptors by two distinct mechanisms. Nature 408, 936–943. (2000).

24. Bats, C., Groc, L. & Choquet, D. The interaction between Stargazin and PSD-95 regulates AMPA receptor surface trafficking. Neuron 53, 719–734 (2007).

25. Cho, C.H., St-Gelais, F., Zhang, W., Tomita, S. & Howe, J.R. Two families of TARP isoforms that have distinct effects on the kinetic properties of AMPA receptors and synaptic currents. Neuron 55, 890–904 (2007).

26. Schwenk, J. et al. Regional diversity and developmental dynamics of the AMPA-receptor proteome in the mammalian brain. Neuron 84, 41–54 (2014).

27. Tomita, S. et al. Functional studies and distribution define a family of transmembrane AMPA receptor regulatory proteins. The Journal of cell biology 161, 805–816 (2003).

28. Milstein, A.D., Zhou, W., Karimzadegan, S., Bredt, D.S. & Nicoll, R.A. TARP subtypes differentially and dose-dependently control synaptic AMPA receptor gating. Neuron 55, 905–918 (2007).

29. Kott, S., Werner, M., Korber, C. & Hollmann, M. Electrophysiological properties of AMPA receptors are differentially modulated depending on the associated member of the TARP family. J Neurosci 27, 3780–3789 (2007).

30. Fukaya, M. et al. Abundant distribution of TARP gamma-8 in synaptic and extrasynaptic surface of hippocampal neurons and its major role in AMPA receptor expression on spines and dendrites. The European journal of neuroscience 24, 2177–2190 (2006).

31. Inamura, M. et al. Differential localization and regulation of stargazin-like protein, gamma-8 and stargazin in the plasma membrane of hippocampal and cortical neurons. Neurosci Res 55, 45–53 (2006).

32. Shaikh, S.A. et al. Stargazin Modulation of AMPA Receptors. Cell Rep 17, 328–335 (2016).

33. Twomey, E.C., Yelshanskaya, M.V., Grassucci, R.A., Frank, J. & Sobolevsky, A.I. Elucidation of AMPA receptor-stargazin complexes by cryo-electron microscopy. Science 353, 83–86 (2016).

34. Zhao, Y., Chen, S., Yoshioka, C., Baconguis, I. & Gouaux, E. Architecture of fully occupied GluA2 AMPA receptor-TARP complex elucidated by cryo-EM. Nature 536, 108–111 (2016).

35. Herguedas, B. et al. Architecture of the heteromeric GluA1/2 AMPA receptor in complex with the auxiliary subunit TARP gamma8. Science 364 (2019).

36. Turetsky, D., Garringer, E. & Patneau, D.K. Stargazin modulates native AMPA receptor functional properties by two distinct mechanisms. J Neurosci 25, 7438–7448 (2005).

37. Tomita, S. et al. Stargazin modulates AMPA receptor gating and trafficking by distinct domains. Nature 435, 1052–1058 (2005).

38. Tomita, S., Fukata, M., Nicoll, R.A. & Bredt, D.S. Dynamic interaction of stargazin-like TARPs with cycling AMPA receptors at synapses. Science 303, 1508–1511 (2004).

39. Yamasaki, M. et al. TARP gamma-2 and gamma-8 Differentially Control AMPAR Density Across Schaffer Collateral/Commissural Synapses in the Hippocampal CA1 Area. J Neurosci 36, 4296–4312 (2016).

40. Fukaya, M., Yamazaki, M., Sakimura, K. & Watanabe, M. Spatial diversity in gene expression for VDCCgamma subunit family in developing and adult mouse brains. Neurosci Res 53, 376–383 (2005).

41. Morimoto-Tomita, M. et al. Autoinactivation of neuronal AMPA receptors via glutamate-regulated TARP interaction. Neuron 61, 101–112 (2009).

42. Neubert, F. et al. Bioorthogonal Click Chemistry Enables Site-specific Fluorescence Labeling of Functional NMDA Receptors for Super-Resolution Imaging. Angew Chem Int Ed Engl 57, 16364–16369 (2018).

43. Plass, T., Milles, S., Koehler, C., Schultz, C. & Lemke, E.A. Genetically encoded copper-free click chemistry. Angew Chem Int Ed Engl 50, 3878–3881 (2011).

44. Serfling, R. et al. Designer tRNAs for efficient incorporation of non-canonical amino acids by the pyrrolysine system in mammalian cells. Nucleic Acids Res 46, 1–10 (2018).

45. Blackman, M.L., Royzen, M. & Fox, J.M. Tetrazine ligation: fast bioconjugation based on inverseelectron-demand Diels-Alder reactivity. J Am Chem Soc 130, 13518–13519 (2008).

46. Yasuda, R. Imaging spatiotemporal dynamics of neuronal signaling using fluorescence resonance energy transfer and fluorescence lifetime imaging microscopy. Curr Opin Neurobiol 16, 551–561 (2006).

47. Priel, A. et al. Stargazin reduces desensitization and slows deactivation of the AMPA-type glutamate receptors. J Neurosci 25, 2682–2686 (2005).

48. Rimbault, C. et al. Engineering selective competitors for the discrimination of highly conserved protein-protein interaction modules. Nat Commun 10, 4521 (2019).

49. Heilemann, M. et al. Subdiffraction-resolution fluorescence imaging with conventional fluorescent probes. Angew Chem Int Ed Engl 47, 6172–6176 (2008).

50. van de Linde, S. et al. Direct stochastic optical reconstruction microscopy with standard fluorescent probes. Nat Protoc 6, 991–1009 (2011).

51. Louros, S.R., Caldeira, G.L. & Carvalho, A.L. Stargazin Dephosphorylation Mediates Homeostatic Synaptic Downscaling of Excitatory Synapses. Front Mol Neurosci 11, 328 (2018).

52. Oliveira, B.L., Guo, Z. & Bernardes, G.J.L. Inverse electron demand Diels-Alder reactions in chemical biology. Chem Soc Rev 46, 4895–4950 (2017).

53. Karver, M.R., Weissleder, R. & Hilderbrand, S.A. Synthesis and evaluation of a series of 1,2,4,5-tetrazines for bioorthogonal conjugation. Bioconjug Chem 22, 2263–2270 (2011).

54. Lajoie, M.J. et al. Genomically recoded organisms expand biological functions. Science 342, 357–360 (2013).

55. Rackham, O. & Chin, J.W. A network of orthogonal ribosome x mRNA pairs. Nat Chem Biol 1, 159–166 (2005).

56. Wang, K., Neumann, H., Peak-Chew, S.Y. & Chin, J.W. Evolved orthogonal ribosomes enhance the efficiency of synthetic genetic code expansion. Nat Biotechnol 25, 770–777 (2007).

57. Neumann, H., Wang, K., Davis, L., Garcia-Alai, M. & Chin, J.W. Encoding multiple unnatural amino acids via evolution of a quadruplet-decoding ribosome. Nature 464, 441–444 (2010).

58. Beliu, G. et al. Tethered agonist exposure in intact adhesion/class B2 GPCRs through intrinsic structural flexibility of the GAIN domain. Mol Cell (2021).

59. Nikic, I. et al. Debugging Eukaryotic Genetic Code Expansion for Site-Specific Click-PAINT SuperResolution Microscopy. Angew Chem Int Ed Engl 55, 16172–16176 (2016).

60. Constals, A. et al. Glutamate-induced AMPA receptor desensitization increases their mobility and modulates short-term plasticity through unbinding from Stargazin. Neuron 85, 787–803 (2015).

61. Kaech, S. & Banker, G. Culturing hippocampal neurons. Nat Protoc 1, 2406–2415 (2006).

62. Stoppini, L., Buchs, P.A. & Muller, D. A simple method for organotypic cultures of nervous tissue. J Neurosci Methods 37, 173–182 (1991).

63. Wolter, S. et al. rapidSTORM: accurate, fast open-source software for localization microscopy. Nat Methods 9, 1040–1041 (2012).

64. Kiskowski, M.A., Hancock, J.F. & Kenworthy, A.K. On the use of Ripley’s K-function and its derivatives to analyze domain size. Biophys J 97, 1095–1103 (2009).

65. Dixon, P.M. in Encyclopedia of Environmetrics (2001).

66. Hafner, A.S. et al. Lengthening of the Stargazin Cytoplasmic Tail Increases Synaptic Transmission by Promoting Interaction to Deeper Domains of PSD-95. Neuron 86, 475–489 (2015).

